# Revisiting the evolution and function of NIP2 parologs in the *Rhynchosporium spp.* complex

**DOI:** 10.1101/2024.10.15.618441

**Authors:** Reynaldi Darma, Daniel S. Yu, Megan A. Outram, Yi-Chang Sung, Erin H. Hill, Daniel Croll, Simon J. Williams, Ben Ovenden, Andrew Milgate, Peter S. Solomon, Megan C. McDonald

**Affiliations:** Research School of Biology, The Australian National University, Canberra, ACT 2601, Australia; Department of Protecting Crops and the Environment, Rothamsted Research, Harpenden, Hertfordshire, AL5 2JQ, U.K; Laboratory of Evolutionary Genetics, Institute of Biology, University of Neuchâtel, Neuchâtel, Switzerland; NSW Department of Primary Industries, Wagga Wagga Agricultural Institute, Wagga Wagga, Australia; School of Biosciences, University of Birmingham, Birmingham, U.K

**Keywords:** *Rhynchosporium commune*, effectors, Necrosis-Inducing Protein, NIP2, paralogs

## Abstract

The fungus *Rhynchosporium commune,* the causal agent of barley scald disease, contains a paralogous effector gene family called *Necrosis-Inducing Protein 2* (*NIP2*) and *NIP2-like protein* (*NLP*). However, the function and full genomic context of these paralogs remains uncharacterised. Here we present a highly contiguous long-read assembly of *R. commune* WAI453. Using this assembly, we show that the duplication of the *NIP2* and *NLP* gene families is distributed throughout the genome and pre-dates the speciation of *R. commune* from its sister species. Some *NIP2* paralogs have subsequently been lost or are absent in the sister species. The diversity of these paralogs was examined from *R. commune* global populations and their expression was analysed during *in planta* and *in vitro* growth to analyse the importance of these genes during infection. The majority of *NIP2* and *NLP* paralogs in WAI453 genome were significantly upregulated during plant infection suggesting that the *NIP2* and *NLP* genes harbour virulence roles. An attempt to further characterise function of NIP2.1 by infiltrating purified protein into barley leaves did not induce necrosis questioning its previously reported role as an inducer of host cell death. Together these results suggest that the *NIP2* effector family does play a role during infection of barley, however the exact function of *NIP2*, like many effectors, remains uncharacterised.

## 1. Introduction

*Rhynchosporium commune* is the causal agent of barley scald disease, which can cause grain yield losses as high as 45% under favourable field conditions (Brown, 1985). In the United Kingdom, scald disease causes £7.2 million in yield and grain quality losses to the barley industry despite reliance on fungicide treatment programs (Paveley et al., 2016). The predominant symptom of scald is large necrotic lesions on affected leaves with dark brown margins (Avrova and Knogge, 2012). Scald also severely reduces grain quality leading to lower grain value (Avrova and Knogge, 2012; Skoropad, 1959; Zhan et al., 2008). Early studies of the *Rhynchosporium* genus described two different species, *Rhynchosporium secali*s which infected barley, rye, and some species of wild grasses, and *Rhynchosporium orthosporum* which infected orchard grass (Goodwin, 2002). However, more recent analyses divided *R. secalis* into three different species; *R. commune* which infects barley and other *Hordeum* spp., *R. secalis* which infects rye and triticale, and *Rhynchosporium agropyri* which infects *Agropyron* spp. (Zaffarano et al., 2011). Later a novel *Rhynchosporium* species was isolated from perennial ryegrass and was named *Rhynchosporium lolii* (King et al., 2013). *R. commune*, *R. secalis*, *and R. agropyri* are classified as the beaked conidia group (BCG) and are also phylogenetically closely related (King et al., 2013; Penselin et al., 2016; Zaffarano et al., 2011). The remaining two species, *R. lolii* and *R. orthosporum* are classified as the cylindrical conidia group (CCG) and form a separate phylogenetic group (King et al., 2013). Among these five species, *R. commune* and *R. secalis* are considered the most damaging diseases with a global distribution, whereas the other species remain minor diseases primarily on pasture grass species (King et al., 2013; Paveley et al., 2016; Zaffarano et al., 2011).

*R. commune* is considered a hemibiotropic pathogen, that has a long asymptomatic period (7-10 days), followed by the rapid appearance of large necrotic lesions (Avrova and Knogge, 2012; Kirsten et al., 2012). This two-stage infection cycle is hypothesised to be driven by different sets of secreted effectors. In the first stages of infection effectors that supress plant immunity are hypothesised to be the main drivers of infection, which is then followed by a second set of necrosis-inducing effectors that damage plant tissues (Lu et al., 2022; Shao et al., 2021). To date, three effectors secreted by *R. commune* have been described following purification from *in vitro* culture filtrate. These proteins were named Necrosis-Inducing Proteins (NIP1, NIP2, and NIP3), as they were shown to induce necrosis when infiltrated into barley leaves (Wevelsiep et al., 1991). NIP1 and NIP3 were both shown to stimulate the host plant plasma membrane H^+^-ATPase, whereas the NIP2 was not found to affect ATPase activity (Wevelsiep et al., 1993). Despite the strong activity observed in these studies, no further work has been conducted to explore the mechanistic basis leading to plant necrosis.

The *NIP* effector gene family has since been expanded significantly based on whole genome re-sequencing studies of *R. commune.* Mohd-Assaad et al. (2019) recently reported the presence of two highly identical *NIP1* paralogs, now named *NIP1A* and *NIP1B*. Interestingly, these genes appear to not only exist as paralogs but also have copy number variation in different fungal isolates (Mohd-Assaad et al., 2019). In a global study, *NIP1A* was found more frequently in isolates when compared to *NIP1B* and isolates containing a functional *NIP1A* were also found to be more virulent than those without *NIP1A*. *NIP1B* had a smaller but significant effect on virulence in isolates that carried two copies of this paralog, suggesting that both effector presence/absence polymorphism and copy number variation can play a role in the virulence in this important pathogen (Mohd-Assaad et al., 2019).

Remarkably, *R. commune* carries 11 described *NIP2* paralogs (*NIP2.1-NIP2.11*), defined by three conserved motifs: a 40-amino acid sequence at the N-terminus, a 15-amino acid sequence containing three conserved amino acids (cysteine, arginine and serine (CRS motif)) in the middle of the protein sequence, and a C-terminal 15-amino acid sequence (Penselin et al., 2016). In addition to these three conserved motifs, these 11 paralogs also have six conserved cysteine residues. The *NIP2.1* gene also contains a unique intron in the 3’ untranslated region (UTR) immediately following a stop codon (Kirsten et al., 2012). The presence of this 3’ UTR intron in the other *NIP2* paralogs remains uncharacterised. Two more distantly related NIP2-Like Proteins (*NLPs*) NLP2 and NLP3 have also been described (Penselin et al., 2016). NLP proteins also share the six conserved cysteine residues found in NIP2 paralogs, but are less conserved in the other NIP2 protein domains, most notably lacking the conserved CRS domain (Penselin et al., 2016).

Apart from NIP2.1 that was reported to induce necrosis on barley leaves (Wevelsiep et al., 1991), only one other paralog, NIP2.6 (*RcSP6*) has been functionally investigated (Penselin et al., 2016). The virulence of *RcSP6* knockout mutant was similar to that of wild-type suggesting this gene does not have a role in inducing necrosis on barley leaves. However, the biomass of the mutant was more abundant than the biomass of the wild-type indicating this gene plays a role in supressing fungal growth during the asymptomatic period of infection (Penselin et al., 2016). Together these studies indicate that NIP2.6 and NIP2.1 have different functions within the *R. commune*-barley interaction. In this paper, we assessed the genomic context and evolutionary history of this expansive effector family in *R. commune*. We did this by generating a high-quality contiguous genome assembly and analysing the diversity of NIP2 paralogs from *R. commune* global populations and its sister species. We also examined the expression of the *NIP2* paralog family *in plant*a, to explore if the *NIP2* paralogs have diversified the timing of their expression during infection. Finally, we used heterologous expression to explore the reported necrosis-inducing activity of NIP2.1.

## 2. Materials and methods

### 2.1. *R. commune* strains, fungal genome sequencing and assembly

*R. commune* Australia_New isolates, including WAI453, were collected by the New South Wales Department of Primary Industries, Australia as part of the Wagga Wagga Agricultural Institute Isolate Collection. For long term storage, spores of WAI453 was resuspended in 25% glycerol and stored at -80°C. WAI453 strain was grown on 2% Lima Bean Agar (filtrate of 125 g/L boiled Lima Bean, 20 g/L agar) and incubated at 18°C in darkness for 10 days. Spores were collected from agar plates and the HMW genomic DNA was extracted according to the methods described on protocols.io (dx.doi.org/10.17504/protocols.io.9g3h3yn). The genomic DNA library prepared from the extracted HWM DNA was generated using a PacBio SMRTBell Template Prep Kit 1.0 SPv3 with BluePippin 15-50 kb size selection. The library was sequencing on a PacBio Sequel machine with sequel sequencing kit v2.1 chemistry at the Ramaciotti Centre (UNSW Sydney, Australia).

Raw PacBio sequencing reads were corrected, trimmed, and *de novo* assembled with Canu v1.5 with the following settings: genomeSize=57m, minReadLength=11000, correctedErrorRate=0.040, and stopOnReadQuality=false (Koren et al., 2017). This raw assembly was further polished with raw PacBio reads to obtain a more accurate assembly. To do this, the raw reads were first aligned to the raw assembly using BLASR v5.1 with the following settings: --hitPolicy randombest --minMatch12 --nCandidates 2 --useQuality true --minReadlength 1000 --minSubreadLength 500 (https://github.com/PacificBiosciences/blasr). The raw assembly was subsequently polished with raw genomic reads using Variantcaller Arrow v2.3.2 with the following settings: -minConfidence 40 --minCoverage 25 -- coverage 100 --minReadScore 0.75 (https://github.com/PacificBiosciences/GenomicConsensus).

Short-read sequencing was performed for other 71 *R. commune* Australian_New isolates at The Australian Genome Research Facility (Westmead, NSW, Australia). The genomic DNA libraries were prepared using Nextera DNA Flex and sequenced for 200 cycles paired-end (100bp PE) sequencing using NovaSeq 6000 Illumina machine. The sequencing reads were assembled using SPAdes v3.13 (Bankevich et al., 2012).

### 2.2. Plant infection, RNA sequencing from *in planta* and *in vitro* growth

Plant infection assays were performed on 3-week-old barley cv. ND5883 grown in controlled environment growth chambers under a 14 hour day/ 10 hour night cycle (20°C day/12°C night) at 85% relative humidity according to McDonald et al. (2018). A concentration of 1×10^6^ spores/ml in 0.02% Tween-20 was sprayed on the top of the 3^rd^ leaves of 3-week-old barley seedlings. Infected seedlings were incubated in darkness at 18°C with 100% humidity for 48 hours before they were transferred to a growth chamber for an additional 6 days incubation under normal plant growth conditions.

For *in vitro* growth, 50 ml of Lima Bean Broth was inoculated with 1×10^6^ spores of WAI453 and incubated in the dark with shaking (120 RPM) at 18°C for 8 days. Fungal mycelia and culture filtrate were separated with Miracloth (Merck). Infected leaves and fungal mycelia were snap frozen in liquid nitrogen and stored at -80°C before RNA extraction. RNA from three biological replicates of infected leaves and fungal mycelium were extracted using Quick-RNA Fungal/Bacteria Miniprep (Zymo Research) following the manufacturer’s protocol. Eluted RNA was treated for genomic DNA contamination using TURBO™ DNAse (Thermo Fisher). High-quality RNA from *in vitro* samples and *in planta* samples were used to construct Illumina stranded mRNA libraries. These libraries were sequenced for single-end sequencing using Nextseq high output 75 cycles sequencing platform with 400M reads output at Biomolecular Resource Facility (The Australian National University). Illumina libraries were multiplexed in different ratios to account for the pure fungal reads in the *in vitro* samples (∼20M reads) compared to the mixed plant and fungal reads in the *in planta* samples (∼110M reads).

### 2.3. RNA reads mapping, genome annotation, and differential gene expression analysis

The quality of RNA sequencing raw reads was assessed using FastQC (v0.11.8) (Andrews, 2010). Low quality reads were removed and adapter sequences trimmed using Trimmomatic v0.33 with the following settings: -phred33 ILLUMINACLIP:TruSeq3-SE.fa:2:30:10 SLIDINGWINDOW:4:15 MINLEN:50 (Bolger et al., 2014). Trimmed reads from all *in vitro* and *in planta* samples were then combined and mapped to the WAI453 genome assembly using STAR v2.7.2a with the following settings: --twopassMode Basic--alignIntronMin 10 --alignIntronMax 300 (Dobin et al., 2013). The mapped reads were assembled into transcripts using StringTie v2.0 using stranded library –rf mode (Pertea et al., 2015). Gene model prediction for the WAI453 genome assembly was performed using two gene model prediction software, BRAKER v2.0 (Brůna et al., 2021) and CodingQuarry v2.0 (Testa et al., 2015). Annotation with CodingQuarry v2.0 was performed using the ‘pathogen stranded mode’ and assembled transcripts were used as evidence for gene model prediction. BRAKER v2.0 gene model annotation was performed with mapped RNA-seq reads as input and using the --fungus parameter for fungal-specific intron prediction. Only the longest isoform from each predicted gene model was retained in the BRAKER output. EVidenceModeler v1.1.1 (Haas et al., 2008) was then used to consolidate predicted gene models from CodingQuarry (evidence weight of 10) and BRAKER (evidence weight of 4) into a final annotation.

For differential gene expression (DGE) analysis, trimmed reads from each *in vitro and in planta* sample were individually mapped to the WAI453 genome using STAR v2.7.2a using the following parameters: --alignIntronMin 10 --alignIntronMax 300 and with the new genome annotation as input (Dobin et al., 2013). The mapped reads were assembled into transcripts using StringTie v2.0 with parameters –rf -e -B -G. Mapped reads overlapping fungal coding regions and read counts were then extracted using the Python script (prepDE.py) (Pertea et al., 2015). EdgeR v3.26.8 (Robinson et al., 2010) was subsequently used to perform DGE analysis. Any predicted gene with less than 3 counts per million (cpm) in more than three individual samples was discarded and the trimmed mean of M-values (TMM) method was used in the normalization process. The glmQLFit function, which uses generalized linear models (GLMs) with quasi-likelihood (QL) F-test to test any group of samples, was used to generate a gene expression dataset including their P-value and the False Discovery Rate (FDR value). The DGE group was generated using the Benjamin-Hochberg adjusted FDR method (Benjamini and Hochberg, 1995) with P-value < 0.01, and 2 LogFC or more. Reads per kilobase per million mapped reads of each gene in each biological treatment was calculated with rpkm function in EdgeR library. Script to perform DGE analysis and RPKM calculation is available in Zenodo (DOI:10.5281/zenodo.13651118).

### 2.4. *NIP2* and *NLP* genes detection from global isolates, Haplotype networks and phylogenetic tree

The presence of *NIP2* and *NLP* genes in Australia_New isolates were analysed with BLAST+ v2.9.0+ (Camacho et al., 2009) using NIP2 and NLP sequences as the query input. Python script, BLASTtoGFF_multiple.py (DOI:10.5281/zenodo.13651118), was then used to parse them into fasta file. The *NIP2* and *NLP* genes from other *R. commune* global isolates and its sister species were extracted following the procedure of Mohd-Assaad et al. (Mohd-Assaad et al., 2019).

To generate the haplotype network for each *NIP2* and *NLP* gene, *NIP2* and *NLP* gene sequences were aligned in the Phylip format. Missing nucleotides in the alignment were denoted with an “X”. The alignments were imported into PopART v1.7 (Leigh and Bryant, 2015) which was used to generate a haplotype network for each *NIP2* and *NLP* gene using minimum spanning network.

To generate a phylogenetic tree from NIP2 and NLP proteins present in the *R. commune* global isolates and its sister species, protein sequences were firstly aligned with the Geneious Alignment in Geneious Prime 2019.1.3 (Kearse et al., 2012) with the following settings: global alignment, blosum 62 cost matrix, gap open penalty 12, gap extension penalty 3, refinement iterations 2. The result was then manually edited to obtain a better alignment. The best amino acid substitution model for the phylogenetic tree was identified from the protein alignment result in the IQTree 2.0 program (Kalyaanamoorthy et al., 2017; Nguyen et al., 2015). A Bayesian phylogenetic tree was created using MrBayes 3.2.6 (Huelsenbeck and Ronquist, 2001) with the following settings: WAG Rate Matrix, gamma rate variation, four gamma categories, 1,000,000 chain length, four heated chains, 0.2 heated chain temp, 10,000 subsampling freq, 100,000 burn-in lengths, and 3,246 random seed. A maximum likelihood tree (ML) was also created with RAxML v8.2.11 (Stamatakis, 2014) with the following settings: GAMMA WAG protein model, rapid bootstrapping and search for best–scoring ML tree algorithm, 10,000 bootstrap replicates, and 1,250 parsimony random seed. Finally, the appearance of the phylogenetic tree was polished using FigTree v1.4.4 (Rambaut, 2018).

### 2.5. RNA isolation and qPCR of *NIP2* genes

Barley cv. ND5883 leaves were inoculated with the WAI453 isolate as described above. Infected leaves were collected from different time points: at 0-dpi (less than one hour after inoculation), 3-dpi, 6-dpi, 9-dpi, and 12-dpi. Three biological replicates for each sample were used in this experiment. Total RNA was extracted from those samples using Quick-RNA Fungal/Bacteria Miniprep (Zymo Research) following the manufacturer’s protocol. Genomic DNA contamination was degraded using TURBO™ DNAse (Thermo Fisher). First-strand cDNA was synthesised using SuperScript IV Reverse Transcriptase (Invitrogen) following the manufacturer’s recommendation. cDNA samples were used for qPCR. Reactions were performed with Fast SYBR™ Green Master Mix (ThermoFisher) in a ViiA 7 Real-Time PCR System, following the Fast SYBR™ Green Master Mix recommendations. For calculating the amplification efficiency, standard curves for each primer set was generated according to the method described by Gardiner et al. (2004). The mean normalized expression (MNE) was used to analyse the expression of *NIP2.1*, *NIP2.3*, and *NIP2.6* relative to β-tubulin expression according to Muller et al. (2002). Primers used in this study are listed in Table S3.

### 2.6. NIP2.1 functional studies

The NIP2.1 protein was produced in *E. coli* using the CyDisCo system as previously described (Yu et al., 2022). To test the *in planta* activity of the purified protein, 10 µM or 160 µM of NIP2.1 was infiltrated with a 1 mL needless syringe into the first leaf of 9-day old barley cultivar Atlas 46 (*Rrs1* and *Rrs2*), Atlas (*Rrs2*) and ND5883 (no known major scald resistance genes). Infiltrated plants were incubated for 9 days in the growth chamber, as described above.

### 2.7. NIP2 structure prediction

Structures of NIP2.1, NIP2.3, and NIP2.6 proteins (without their signal peptides) were predicted using Google DeepMind’s AlphaFold colab notebook using default settings (https://colab.research.google.com/github/sokrypton/ColabFold/blob/main/AlphaFold 2.ipynb) (Mirdita et al., 2022). To investigate potential structure-informed biological functions, proteins with a similar structure to the predicted structure of NIP2.1 were searched for using the Foldseek database in TM-align mode (van Kempen et al., 2023) and the DALI server (Holm and Rosenstrom, 2010). The predicted NIP2.1 structure was aligned with other structural similar proteins in the pairwise alignment mode in DALI server (Holm and Rosenstrom, 2010). The predicted structures, and structural alignments, were visualized using PyMOL (The PyMOL Molecular Graphics System, Version 2.5.3 Schrödinger, LLC).

## 3. Results

### 3.1. Generating a highly contiguous genome assembly and annotation for *R. commune* WAI453

To generate a chromosome-scale genome assembly, high-molecular weight (HMW) genomic DNA of *R. commune* WAI453 was sequenced with PacBio Sequel using two SMRT cells, yielding 13.4 Gb of reads (Table S1). The resulting WAI453 assembly contained 23 contigs, of which 20 contigs were nuclear DNA, one contig was mitochondrial DNA, and two contigs contained solely ribosomal repeats (Figure 1). The 20 nuclear contigs had a total length of 57.76 Mbases and the genome N50 was 3.6 Mbases (Table 1). Eleven contigs contained telomeric repeats on at least one end and two contigs had telomeres at their both ends, indicating two full chromosomes were obtained in this assembly. Compared to the previously published *R. commune* reference UK7, the WAI453 genome assembly was 1.04% longer and contained 24X fewer contigs, indicating a significant improvement in the assembly contiguity for this species (Penselin et al., 2016) (Table 1). Genome completeness was assessed with the Benchmarking Universal Single-Copy Orthologs (BUSCO) tool (Simao et al., 2015). Among the 1,315 orthologs present in the BUSCO Ascomycete set, 1299 (98.7%) were present in single copy in the WAI453 assembly (Figure S1).

**Figure 1.**
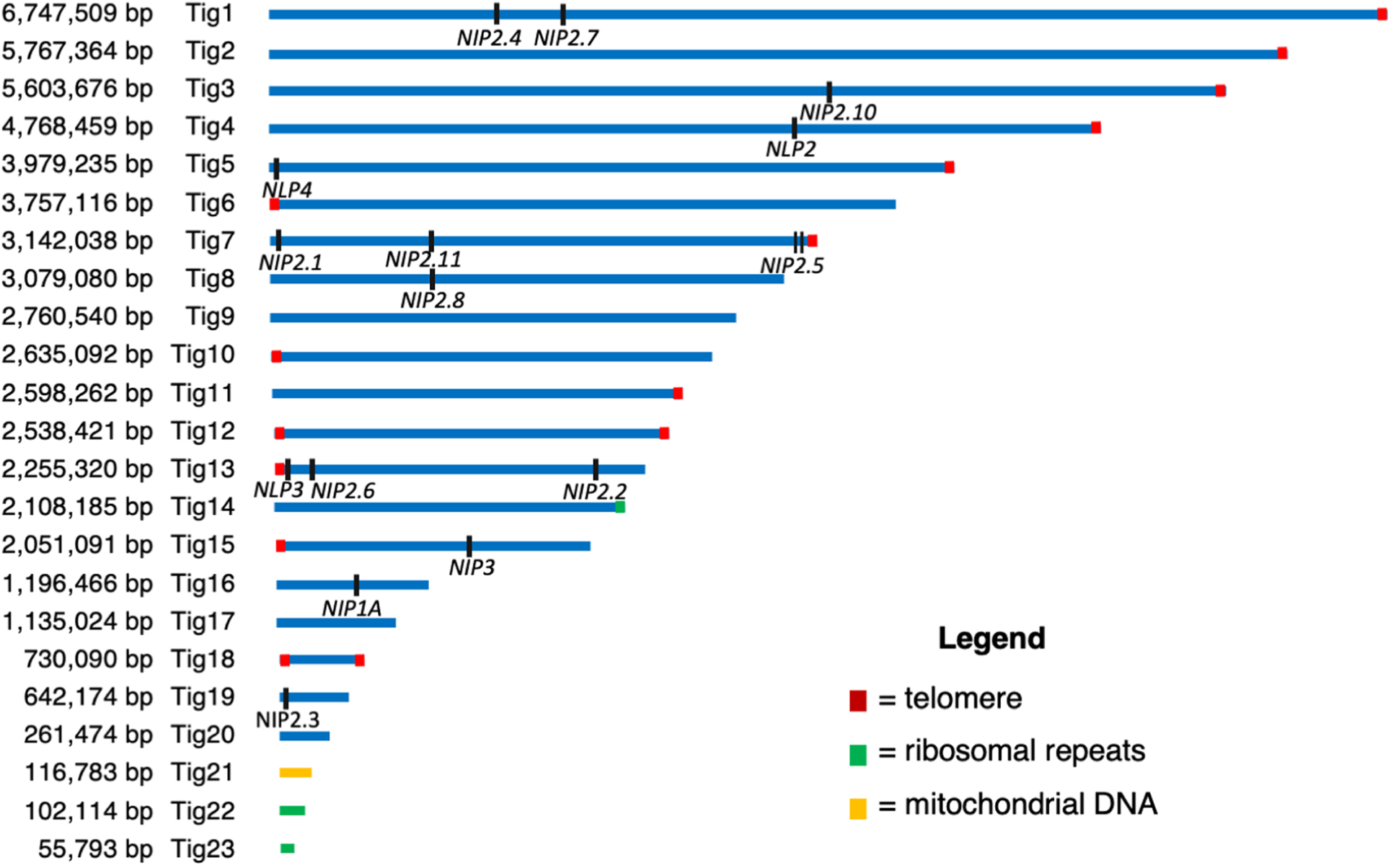
A schematic representation of the *R. commune* WAI453 *de novo* assembly showing all 23 contigs obtained. Nuclear chromosomes are shown as blue lines, whereas mitochondrial and rRNA repeat containing contigs are colored yellow and green, respectively. Contigs that contain telomeric repeats are noted with red boxes. The locations of *NIP1*, *NIP3*, *NIP2* and *NLP* genes in the *R. commune* WAI453 genome assembly are indicated with small black dashes.

**Table 1.** Comparison of *R. commune* WAI453 and *R. commune* UK7 genome assemblies.

| Parameter | WAI453 (PacBio long-read sequencing) | UK7 (Illumina short-read sequencing) (Penselin et al.[9]) |
| --- | --- | --- |
| Coverage | 232x | 267x |
| Genome size | 57.76 Mbases | 55.59 Mbases |
| Contigs | 20 | 481 |
| Scaffolds | - | 163 |
| N50 contig | 3,757 kb | 49.6 kb |
| N50 scaffold | - | 800.5 kb |
| Predicted genes | 13,726<sup>b</sup> | 12,211 |
| No. of predicted secreted Effectors<sup>a</sup> | 445 | 359 |
| Coding regions | 17,158,241 | nd |
| AT-rich region<sup>b</sup> | 30.5% | 29.6% |

<sup>a</sup> analysed by SignalP 6.0 followed by EffectorP 3.0<sup>b</sup> was obtained after *R. commune* WAI453 genome annotation was performed

A genome annotation for this new assembly was also generated using RNA-seq reads obtained from the third leaves of barley (cv. ND5883) 8-days post infection (dpi) and from a 8-day old *in vitro* culture. This annotation contained 13,726 predicted genes and more than 97% of Ascomycete universal single-copy orthologous genes were present (Table 1, Figure S2). Approximately 9,669 predicted proteins were small proteins (<500 amino acids in size), and among these, 445 were predicted as secreted effectors (Figure S3, Table 1). The new annotation was also quality assessed by checking for the correct annotation of the known *NIP2* and *NLP* effector genes. Most of the *NIP2* and *NLP* genes contain an intron in the 3’ untranslated region (UTR) immediately following the stop-codon. Using the RNA-seq data mapped to the WAI453 assembly, the presence of this 3’-UTR-intron was confirmed in the annotation of all *NIP2* and *NLP* genes with one exception in *NIP2.2* (Figure S4). *NIP2.2* is the only paralog that contains an intron in the protein coding sequence which results in a slight shift in the total length of the protein. This paralog is also the only *NIP2* that has a C-terminal extension region after the final cysteine, which was reported previously by Penselin et al. (Penselin et al., 2016).

BLASTn was used to identify the presence and location of all *NIP* and *NLP* genes and the new annotation was cross-referenced with the published sequences for all named effectors. These analyses showed WAI453 carried *NIP1A* but not the paralog *NIP1B.* WAI453 also carried ten *NIP2* genes, missing only *NIP2.9* (Figure 1). In WAI453, three *NIP2* paralogs, *NIP2.5*, *NIP2.10*, and *NLP2*, were determined to be pseudogenes. Among the *NLP* genes, *NLP2 and NL3* were found in this isolate plus one newly described *NLP,* now named *NLP4* (Figure 1). *NLP4* has the same characteristics as other described *NLP genes*, namely the six conserved cysteine residues and the 3’ UTR intron that are also shared with the *NIP2* genes. Together this assembly and annotation gave us a comprehensive view of the entire *NIP2* and *NLP* gene family, which we used for further population genetic and functional analyses.

### 3.2. Exploring the presence of *NIP2* and *NLP* paralogs in a global population and sister species

To comprehensively explore the diversity of all *NIP2* and *NLP* paralogs, we used BLASTn to extract these genes from a global population collection of *de novo* assemblies, analysed previously for the diversity of *NIP1* (Mohd-Assaad et al., 2019) (Table 2). We also added 72 new Australian *R. commune* isolates, including the WAI453, to this dataset. These results showed that with the exception of *NIP2.9,* most *NIP2* genes were largely present in different populations from around the world. *NIP2.9* was the only paralog absent in 86.32% of all isolates globally. The Ethiopian population stood out from other populations as no isolates were found to carry *NIP2.6, NIP2.9* or *NIP2.10* and only partial fragments of *NIP2.11* were identified. This indicates this population has undergone selective loss of these effectors when compared to other regions in the world. No other strong trends were observed that differentiated populations based on the presence or absence of these effectors. Looking further at individual genes, the number of partial gene sequences obtained for *NIP2.5, NIP2.10*, *NIP2.11* and *NLP2* were much higher when compared to other paralogs (Figure 2A, Figure S5). In particular, *NIP2.10* and *NLP2* had a high proportion (>57%) of isolates carrying haplotypes with a premature stop codon (Figure S5). In contrast, all *R. commune* isolates in the global population had the complete *NIP2.4* and *NIP2.7* genes while the *NIP2.3*, *NIP2.8*, and *NLP3* genes were present in 99.47% of isolates (Figure 2A). The majority of the *R. commune* isolates in the global population also carried complete *NIP2.1* (95.79%)*, NIP2.2* (94.74), and *NLP4* genes (97.37%). *NIP2.3* was the most highly conserved effector with a single nucleotide haplotype found in all 189 isolates that had a complete *NIP2.3* gene (Figure S5). Whereas *NIP2.1,* proposed to have necrosis activity, had the highest number of nucleotide haplotypes (N=11) when compared to all other paralogs (Figure S5).

**Figure 2.**
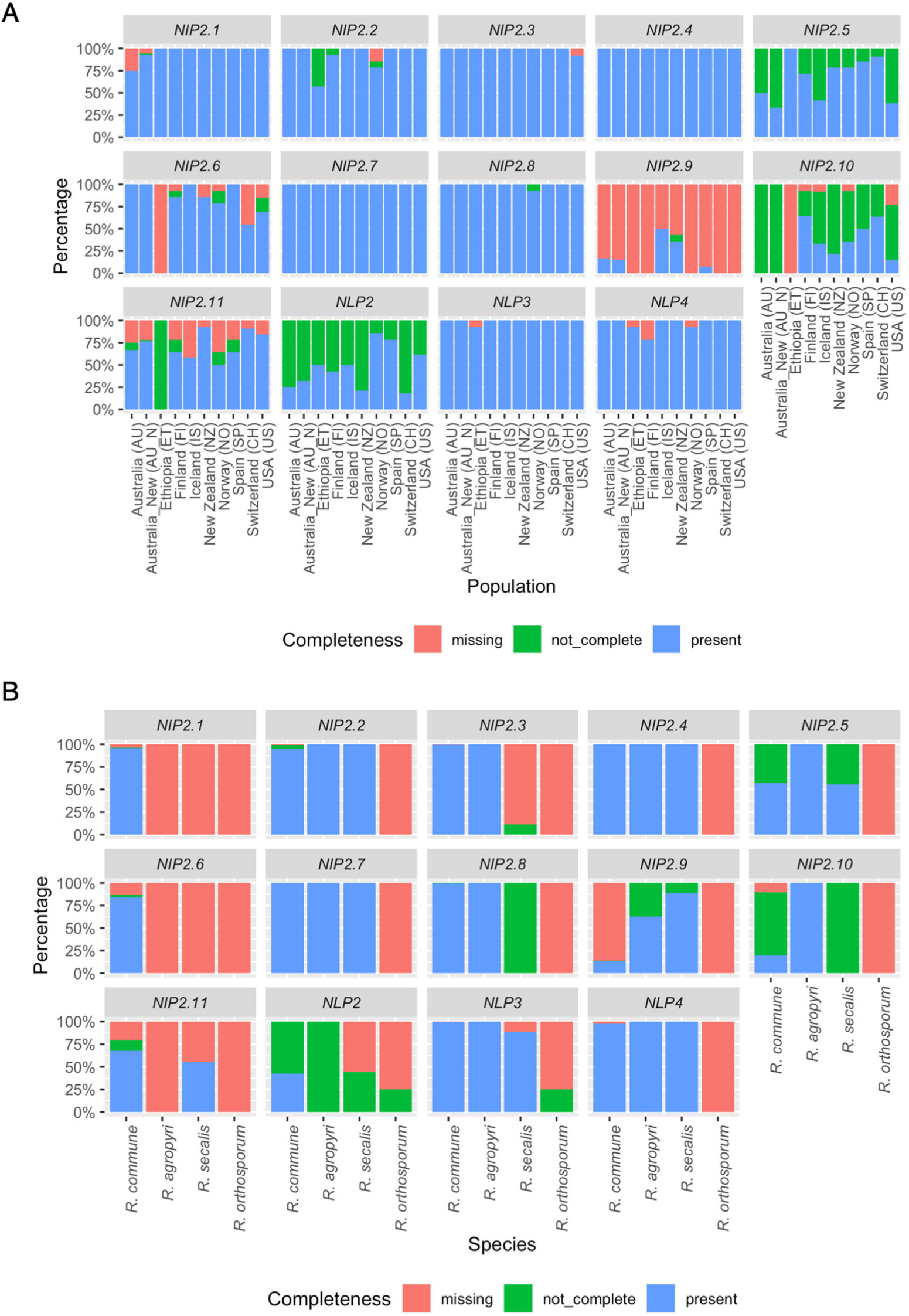
The presence-absence polymorphism of all *NIP2* and *NLP* genes in (A) global *R. commune* isolates, and (B) *R. commune* and three *R. commune* sister species. The presence of these genes are categorised into three groups, which are: Present – isolate had a complete gene, not_complete – isolate only had some parts of the gene or had a premature stop codon inside the gene, and missing – isolate did not have the gene.

**Table 2.**
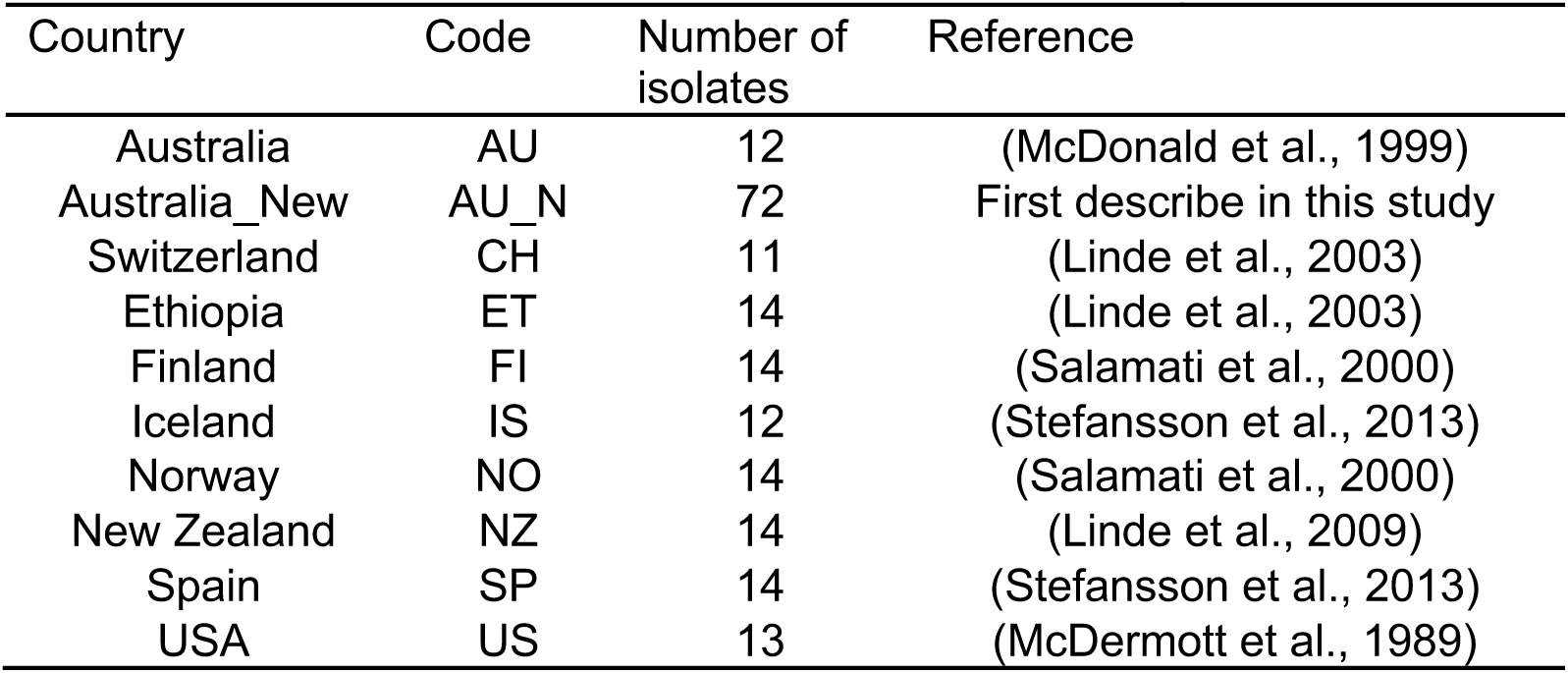
List of *R. commune* isolates used in this study.

The presence of *NIP2* and *NLP* genes were also assessed in the three sister species of *R. commune*, *R. agropyri, R. secalis*, and the more distantly related *R. orthosporum* (Table S2). All *R. agropyri* and *R. secalis* isolates had either complete or partial copies of *NIP2.2-5, NIP2.7-10*, *NLP3 and NLP4* (Figure 2B). On the other hand, *NIP2.1* and *NIP2.6* were only present in *R. commune.* The most distantly related species, *R. orthosporum*, did not carry any of the eleven *NIP2* paralogs and only partial sequences of *NLP2* and *NLP3* in one out of four sequenced isolates. To better understand the evolution of this gene family within the species complex, a phylogenetic tree was constructed using all complete NIP2 and NLP proteins from *R. commune* and its sister species. The amino acid alignment of these NIP2 and NLP protein sequences showed the six conserved cysteine residues, while only NIP2 sequences but not NLP sequences had the CRS domain in the 66-68 residues in the alignment (Figure S6). In this phylogenetic tree, each numbered NIP2 and NLP paralog grouped together with its sister species indicating that gene duplication occurred before speciation (Figure 3). Given the absence of all *NIP2* genes in *R. orthosporum* but presence of *NLP* gene fragments, it appears that the *NIP2* gene expansion occurred after the separation of the beaked conidial group species from the cylindrical conidial species.

**Figure 3.**
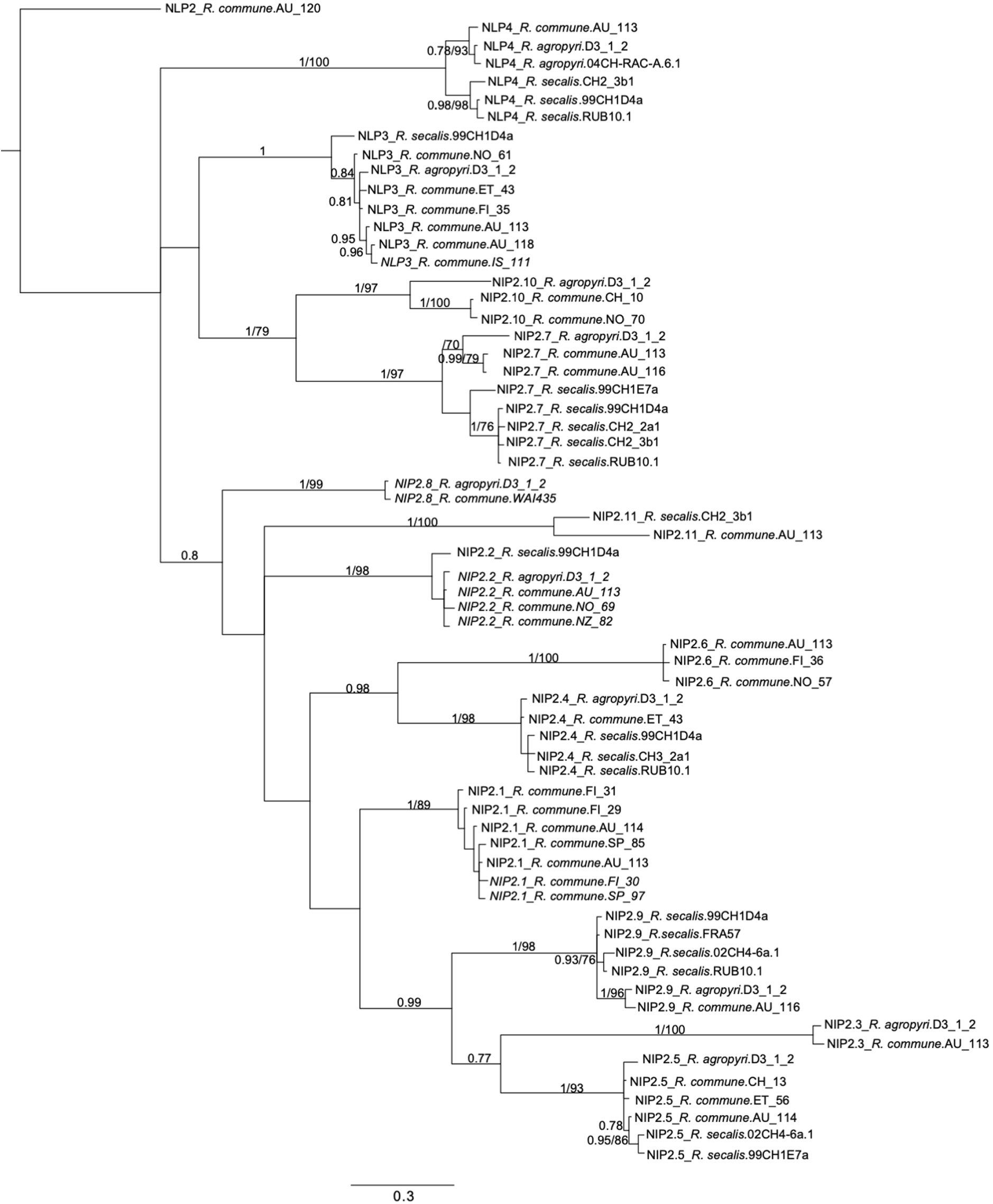
Bayesian inference of phylogenetic tree of mature (without signal peptide) and complete (full length/without premature stop codon) NIP2 and NLP proteins from *R. commune* and its sister species. Numbers on the branches represent Bayesian posterior probabilities (≥0.70)/RAxML bootstrap (≥70%) support values. Unique NIP2 sequences from *R. agropyri* 04CH-RAC-A.6.1 and *R. secalis* 02CH4-6a.1 were also used in this analysis (Penselin et al., 2016).

### 3.3. *NIP2* and *NLP* genes are upregulated during *in planta* growth

Our global screen indicated that most of *NIP2* and *NLP* genes are present in *R. commune* global isolates and its sister species. However, there were distinct differences in sequence conservation between some paralogs (*NIP2.3* only one sequence haplotype globally) compared to paralogs which appear to be pseudogenes in a large number of isolates (*NIP2.5, NIP2.10,* and *NLP2*). To further analyse how important these genes for *R. commune* during infection, the expression of these genes in WAI453 was compared between 8-dpi infected barley (*in planta,* IP samples) and 8-day old *in vitro* culture (IV samples). A multidimensional scaling (MDS) plot was created from total aligned reads of all samples to compare the overall transcriptome profile of *in vitro* vs. *in planta* samples (Figure 4A). This analysis showed all biological replicates from each group were tightly clustered, indicating a similar expression profile between the three biological replicates for each treatment. This analysis also showed, as expected, a clear separation between both growth conditions (Figure 4A). Differential gene expression analysis showed there were 1061 up-regulated genes *in planta* (FDR 5%, P-value< 0.01), while only 418 genes were down-regulated (Figure 4B). The expression of *NIP2* and *NLP* genes was then analysed from the DGE dataset. From 10 fully coding *NIP2* and *NLP* genes present in WAI453 genome, we found seven were significantly upregulated during *in planta* infection, ranging from 2.08 - 6.93 log fold change (LogFC) (Table 3). Neither *NIP2.8* and *NIP2.11* were expressed under the growth conditions tested in this study. The expression of *NIP2.2*, *NIP2.3*, *NIP2.4, NIP2.6*, and *NIP2.7* was low during *in vitro* growth (<35 RPKM), but was significantly induced *in planta* (90.86-324.07 RPKM) (Table 3). In contrast, *NIP2.1* and *NLP3* were highly expressed in both growth conditions. *NLP3* had the highest expression level during *in planta* growth with almost 5,000 RPKM, while its *in vitro* expression peaked at 115 RPKM. *NIP2.1* was not significantly differentially expressed (1.87 LogFC), but this gene was the second most highly expressed during the onset of necrosis with more than 800 RPKM *in planta* and 219 RPKM *in vitro* (Table 3). The newly described *NLP4* was significantly differentially expressed between the two growth conditions, but overall very lowly expressed when compared to the other paralogs.

**Figure 4.**
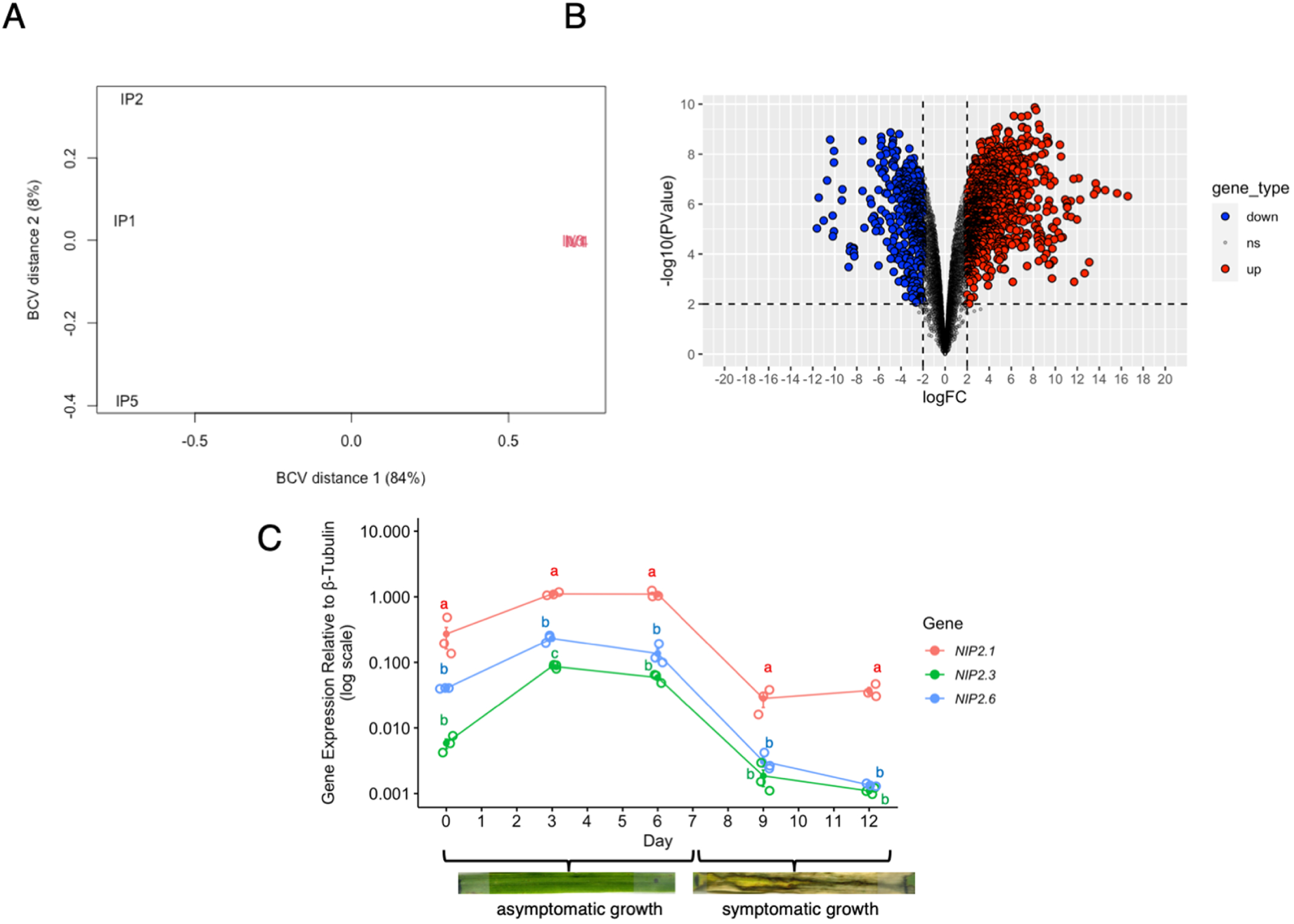
(A) Multidimensional scaling (MDS) plot of the gene expression of *R. commune* WAI453 at 8-dpi on barley cv. ND5883 (labelled as IP) versus *in vitro* culture at 8 days (labelled as IV). The first biological coefficient of variation (BCV distance 1) separates IP and IV. (B) Vulcano plot of the differentially expressed genes. Genes were considered differentially expressed as their log fold change (LogFC) were more than 2 or less than -2 and the P-value less than 0.01. There were 1061 up-regulated genes (2 LogFC or more), 418 down-regulated genes (-2 LogFC or less), and 8144 not differentially expressed genes in the 8-dpi *in planta* growth. Red: up-regulated genes. Blue: down-regulated genes. (C) Gene expression of *NIP2.1*, *NIP2.3*, and *NIP2.6* in the *R. commune* WAI453 during different time points during infection, including asymptomatic and symptomatic growth. The necrotrophic switch was observed at 7-dpi. The expression of *NIP2* genes was monitored by real time quantitative PCR (RT-qPCR) and normalized to *β-tubulin* expression. Standard error of the mean of three biological replicates (RNA from three separate leaves) is shown for each gene and time point. Data point for each replicate was shown using “jitter” function in gglot2. One-way ANOVA with least significant difference (LSD) was performed to analyse the difference between *NIP2* genes expression in each time point. The same letter in each time point suggests there are not significantly different at P-value < 0.05.

**Table 3.**
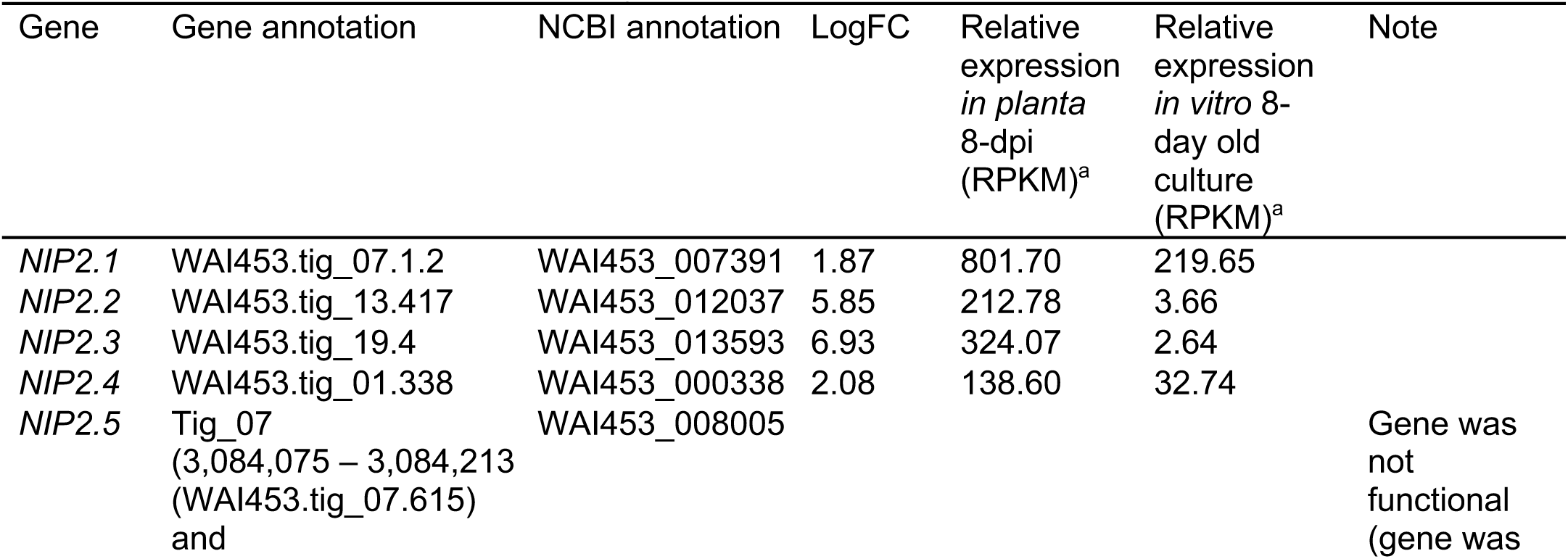
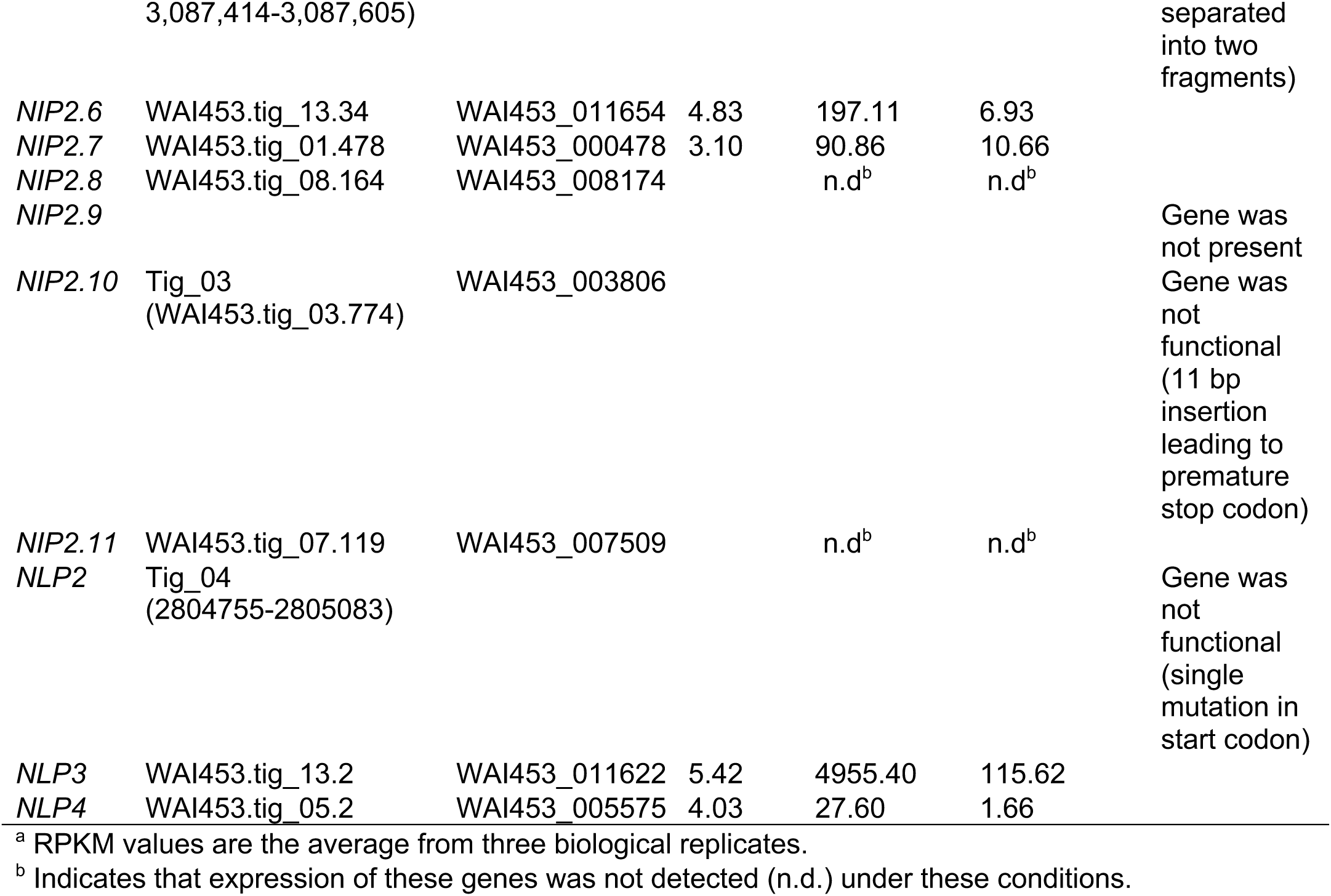
Transcript levels of *NIP2* and *NLP* genes in barley cv. ND5883 leaves infected with *R. commune WAI453 at 8-dpi and in 8-day old R. commune WAI453 in vitro culture*.

The DGE analysis was only performed at a single time point *in planta* (8-dpi), which did not give information regarding of the trajectory of the expression of each of the *NIP2* genes throughout infection. Therefore, we performed qPCR on the *NIP2* genes at 0-dpi (less than one hour after inoculation), 3-dpi, 6-dpi, 9-dpi, and 12-dpi to assess their expression during both the latent and necrotrophic phases of infection. In this experiment, the onset of necrosis was at 7-dpi. Given the large number of time points, this analysis was also limited to three phylogenetically distant *NIP2* genes, *NIP2.1, NIP2.3*, and *NIP2.6* (Figure 3). The expression of *NIP2.1* was significantly higher than the expression of *NIP2.3* or *NIP2.6* in all time points tested, including the 0-dpi time point (Figure 4C). The gene expression of these three genes at 0-dpi was similar with what we observed in the RNA-seq analysis with *in vitro* samples where *NIP2.3* and *NIP2.6* were not expressed or lowly expressed but the *NIP2.1* expression reached 219 RPKM (Table 3). Over the course of these five time points, all three *NIP2* genes were most highly expressed during asymptomatic growth (3-dpi) and immediately prior to the onset of necrosis (6-dpi) (Figure 4C). *NIP2.1, NIP2.3*, and *NIP2.6* were downregulated during the later stages of infection (post 9-dpi) (Figure 4C). Peak expression of *NIP2.1* was observed at the early stage of infection, which raised the question of whether or not *NIP2.1* had a function in inducing plant cell death.

### 3.4. NIP2.1 protein is not a broad-spectrum necrosis-inducing protein in barley

NIP2 was originally described as one of three necrosis-inducing peptides (NIP1, 2 and 3) isolated from *R. commune* culture filtrates (Wevelsiep et al., 1991). According to Wevelsiep and colleagues (1991), NIP2 represented a 6.8 kDa non-glycosylated secreted protein. The protein was identified in culture filtrates in all seven *R. commune* races tested. NIP2 protein purified from culture filtrate of the US238.1 strain caused necrosis at protein concentrations down to ∼50µg/mL (∼7µM) in barley cultivars Atlas and Atlas 46. In a subsequent study, Kirsten and colleagues (2012) cloned the apparent NIP2 protein utilising N-terminal sequence data, referencing Wevelsiep et al. (1991), although these sequencing data were not reported in that study. The *NIP2* gene was subsequently cloned using a PCR walking strategy, with the cloned gene encoding a 93 amino-acid mature protein (without signal peptide) with a molecular weight of ∼10 kDa, as verified by mass spectrometry (MS), which differs from the original molecular weight of NIP2 (Kirsten et al., 2012). The discrepancies in the reported molecular weight of NIP2 has not been addressed.

While studies since Wevelsiep et al., (1991) have confirmed the necrosis activity of NIP1 (Fiegen and Knogge, 2002), to the best of our knowledge no independent study has confirmed the necrosis activity of NIP2, despite the reported difference in size of the protein between the two studies described above. To address this, we heterologously expressed NIP2.1 in *Escherichia coli* strain SHuffle® (Lobstein et al., 2016) with CyDisCo (cytoplasmic disulfide bond formation in the *E. coli*) system (Hatahet et al., 2010; Matos et al., 2014; Yu et al., 2022). We have used this system to investigate other necrosis inducing proteins including Tox1 and Tox3 from *Parastagonospora nodorum* (Outram et al., 2021; Zhang et al., 2016). NIP2.1 protein was purified to homogeneity and underwent quality control experiments, including the use of circular dichroism to demonstrate the protein was correctly folded, and intact protein MS to verify the formation of disulfide bonds, prior to infiltration experiments (Yu et al., 2022).

Pure, folded, NIP2.1 protein was infiltrated into three different barley cultivars at concentrations 10 μM or 160 μM, which were similar to and above that used by Wevelsiep et al. (1991) (estimated to be ∼7-20μM). The three barley cultivars used were Atlas, which has the single dominant scald resistance gene *Rrs2*, Atlas 46, which has both *Rrs1* and *Rrs2* resistance genes, and ND5883, which has no known major scald resistance genes (Goodwin et al., 1990; Wallwork and Grcic, 2011; Zhang et al., 2020). We did not observe necrosis on barley cultivars tested at any of the NIP2.1 concentrations at 9-dpi (Figure 5). Collectively, these data question the role of NIP2.1 as a necrosis inducing protein.

**Figure 5.**
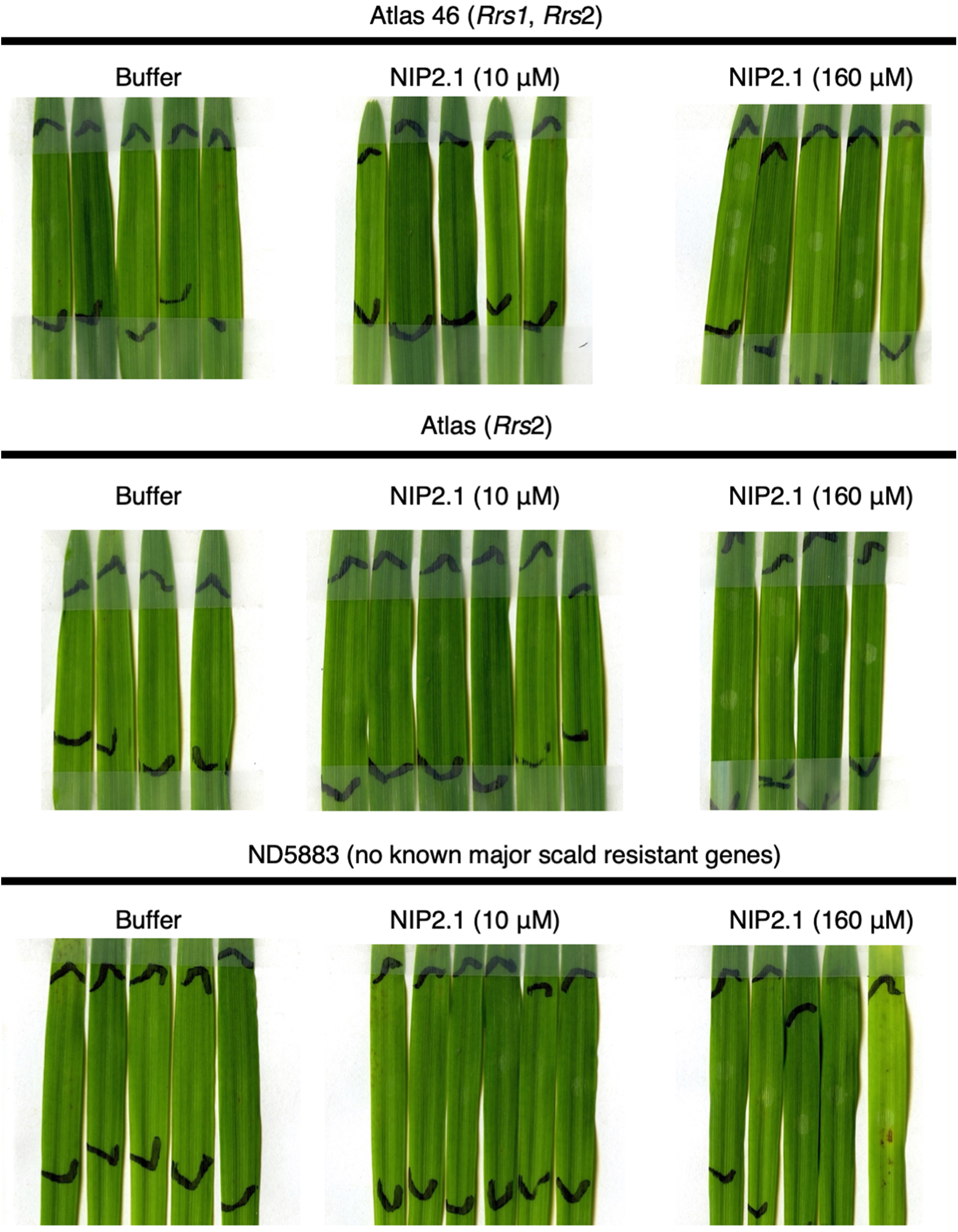
Leaf infiltration assay of pure heterologously produced NIP2.1 protein. NIP2.1 protein at two concentrations (10 μM and 160 μM) was infiltrated into the first leaf of 9-day old barley cultivar Atlas 46 (*Rrs1* and *Rrs2*), Atlas (*Rrs2*) and ND5883 (no known major scald resistance genes). Images were taken at 9 days post infiltration. Black lines indicate the infiltration boundaries.

Like many effectors, the sequence of NIP2 paralogs provide little insight concerning its potential function and role in pathogen virulence. We subsequently sought to utilise recent advances in AI-based protein structure prediction in the form of AlphaFold (Jumper et al., 2021; Mirdita et al., 2022) to predict the structure of NIP2 and investigate for structure-based similarities. The AlphaFold 2–generated NIP2.1 structure was predicted with high confidence (average predicted local distance difference test (pLDDT) score of ∼90 (out of 100) (Figure 6A). The predicted structure consists of five stranded beta-sheet with a discontinuous alpha-helix connecting the β-1 and 2 strands. The model includes three disulfide bonds, which we demonstrated experimentally (Yu et al., 2022). The AlphaFold 2–generated NIP2.3 and NIP2.6 structures were also predicted with high confidence with pLDDT ∼89 and ∼86, respectively (Figure 6A). One of the disulfides was not predicted to form in these NIP2.3 and NIP2.6 models but the cysteines localise to the same region in the NIP2.1 model and therefore likely form a disulphide as in the NIP2.1 predicted structure (Figure S7). This analysis suggests NIP2 paralogs potentially have a similar protein structure.

**Figure 6.**
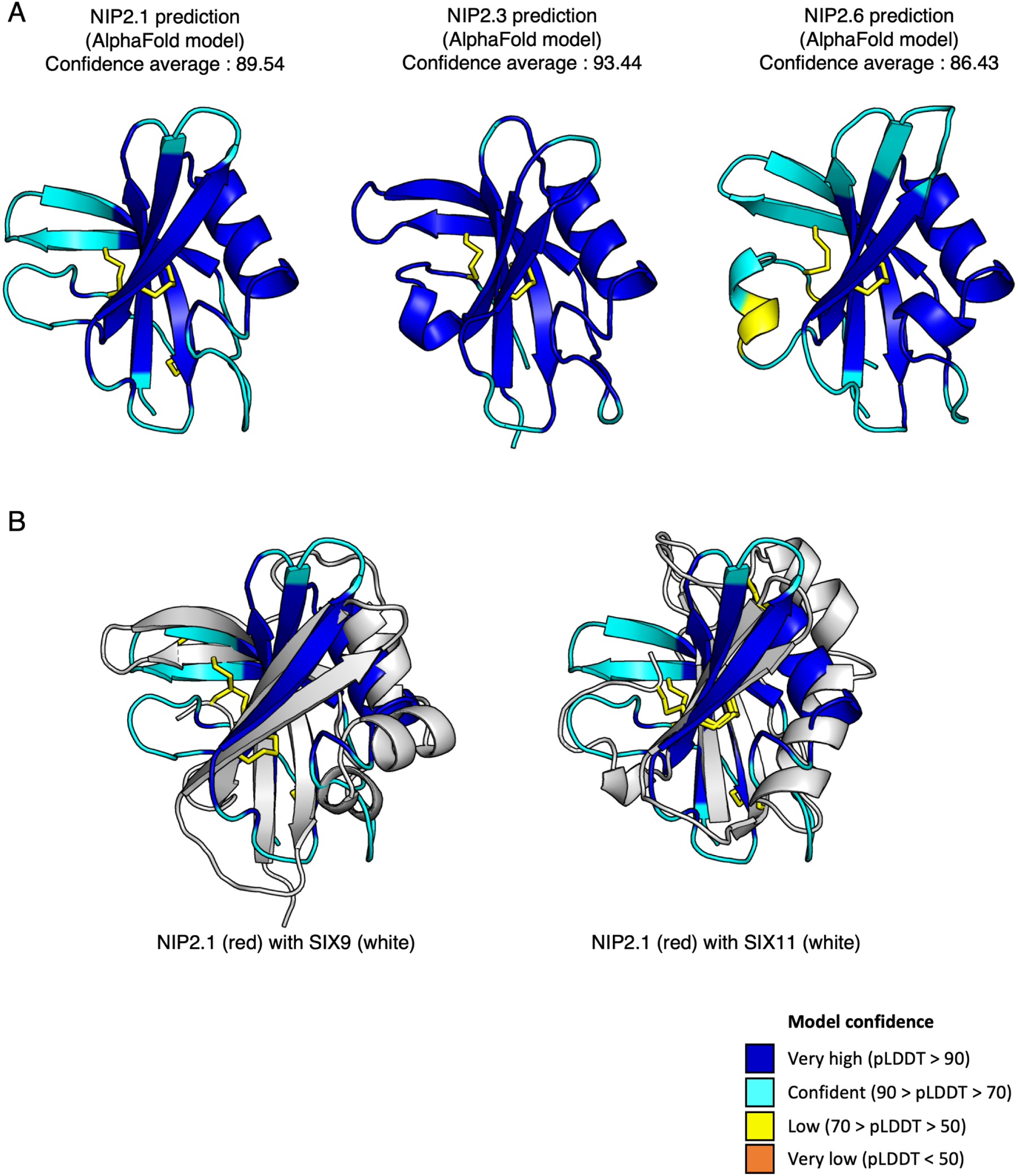
(A) Structural prediction of mature (without signal peptide) NIP2.1, NIP2.3, and NIP2.6 proteins generated by AlphaFold 2. Each residue of each protein was colored based on pLDDT confidence score of AlphaFold 2 prediction. The predicted disulphide bridges are visualised in yellow sticks. (B) NIP2.1 predicted structure aligned with AlphaFold models of family 3 SIX effectors (Yu et al., 2024) in white. SIX9 (left) and SIX11 (right) superimpose with an RSMD of 2.5 and 3.7, respectively (Holm and Rosenstrom, 2010).

Comparison of the NIP2.1 model with experimentally derived structures, utilising the Dali server (Holm and Rosenstrom, 2010), provided little potential functional insights. A broader comparison of the predicted AlphaFold structural database, utilising the Foldseek server (van Kempen et al., 2023), suggested that NIP2.1 shares strong structural similarities with numerous putative fungal effector proteins from other *Rhynchosporium*, *Fusarium* and *Colletotrichum* genus, and known effectors include the *Fusarium oxysporum* secreted-in-xylem (SIX) effectors *SIX9* and *SIX11* (Yu et al., 2024) with TM-score ranging from 0.559 to 0.838. Further analysis by superimposing the predicted NIP2.1 structure with either AlphaFold model of SIX9 or SIX11 structure showed their structures were similar with an RSMD of 2.5 and 3.7, respectively (Figure 6B). Taken together, by combining functional analysis and computational studies, our results suggest that NIP2 protein, at least under the conditions used in our assays, does not induce necrosis on barley leaves.

## 4. Discussion

Here, we described the presence and diversity of the *NIP2* and *NLP* genes across the *Rhynchosporium* spp. complex. This work shows that the majority of *NIP2* and *NLP* genes were present in the global population of *R. commune* and its sister species *R. agropyri and R. secalis* suggesting their importance for adaptability in different hosts and environments. Many *NIP2* genes were highly expressed during *in planta* growth suggesting a role in facilitating infection. However, the NIP2.1 protein, following infiltration, did not induce necrosis on the barley leaves, questioning the original hypothesis of its role in inducing cell death (Wevelsiep et al., 1991). Thus, the RNA-sequencing data and functional protein analyses suggest that *NIP2* genes were important for colonization and infection, although the precise mechanism by which these genes facilitate infection remains unknown.

The duplication of *NIP2* and *NLP* genes was predicted to be an ancient duplication. This is supported by both the assembled genomic location of each paralog and inter-specific phylogenetic analyses conducted in this study. The *de novo* assembly generated for WAI453 is now the most contiguous and complete assembly available for this species. The total size of the assembly falls within the expected range of this species at ∼57 Mb. When examining the distribution of the *NIP2* and *NLP* genes, there was very little evidence of clustering of these genes, with most copies found on unique contigs and or several thousands of kilobases apart from each other. In previous work Mohd-Assaad et al. (2019) noted that extra copies of *NIP1A* and *NIP1B* were located on smaller than average scaffolds, and proposed that these highly identical copies may have been generated by tandem duplications during non-allelic homologous recombination. In WAI453 only a single copy of *NIP1A* was detected, so we were unable to explore this hypothesis further using our long-read sequencing data.

Each numbered paralog grouped more closely with the same paralog from a different species, indicating that these gene duplication events preceded speciation events. Of the eleven described *NIP2* genes, only *NIP2.6* was found in *R. commune* but not in its sister species. While this gene forms a monophyletic group with *NIP2.4*, it remains unclear if this gene arose in *R. commune* alone or instead it has been lost in the other two sister species. Given the small sample size of available genomes for *R. secalis* (N=9) and *R. agropyri* (N=8), it also is likely this gene was simply not present in the available sequenced genomes. Similarly, no *NIP2* genes were detected in the four available *R. orthosporum* genomes. However, there was evidence of partial *NLP* gene sequences. Penselin et al. (2016) showed that *NIP2.1* was also present in other *Rhynchosporium* species, including *R*. *orthosporum* and *R. lolii*, indicating that, at least *NIP2.1* and or *NLP* genes were present before the separation of the *Rhynchosporium* BCG and CCG and may serve as the origin of these paralogs.

In pathogenic fungi, gene duplication and subsequent sequence divergence could lead to neofunctionalization of those duplicated genes (Seong and Krasileva, 2023; Shen et al., 2013). For example, the *Ustilago maydis* effectors *Tay1* and *Mer1* share 31% protein sequence identity and have comparable functions to suppress the reactive oxygen species (ROS) burst, but have different host targets (Navarrete et al., 2021). Tay1 acts in the cytoplasm while the Mer1 acts in the nucleus suggesting they have a different mode of action. Another example of gene duplication leading to neofunctionalisation is the expanded crinkling and necrosis inducing proteins (CRN) of *Phytophthora sojae*, which have less than 50% sequence similarity between half of those gene members, display diverse biological functions such as the induction of apoptosis or a contrasting role in the suppression of programmed cell death (Shen et al., 2013). In addition, Avr4 of *Pseudocercospora fuligena* and its paralog, Avr4-2, are another example of neofunctionalization after gene duplication in pathogenic fungi. These proteins share 33% sequence identity but have different modes how to interact with the plant host (Chen et al., 2021). PfAvr4 binds chitin and protects the pathogen from chitinases whereas PfAvr4-2 binds to de-esterified pectin of primary cell walls or the middle lamella of plant cells to disrupt the cell wall formation (Chen et al., 2021). Gene duplication, however, can also lead to relaxed selection, where secondary copies simply become inactive (Lynch and Conery, 2000; Seong and Krasileva, 2023). There is some evidence of inactivation of some NIP2 paralogs, as over 57% of isolates found carrying *NIP2.10* and *NLP2* carry haplotypes that contain early stop-codons. Similar to other paralogous effectors, the divergence of sequence from NIP2 and NLP proteins leads to the hypothesis that some of them might undergo neofunctionalization or nonfunctionalization, especially for those paralogs that have a different gene expression pattern or those paralogs that mostly absent or present as incomplete proteins in the global population.

In this study, we found that some of the *NIP2* and *NLP* genes were found in most *R. commune* isolates, and many of those paralogs were highly expressed in WAI453 during infection. These findings suggest that the presence of multiple *NIP2* and *NLP* genes were important for *R. commune* during *in planta* growth. The presence of multiple copies of paralogs can be an important feature in fungal pathogen genomes to escape the plant’s defense and maintain fungal virulence (Ridout et al., 2006). Only one paralog of a fungal effector is usually recognised by a specific plant R protein. For example, the AVRa7-1 (CSEP0059) is recognised by Mla7 while the paralog CSEP0060 is not (Saur et al., 2019). Another example is the two AVR-Pik copies (AVR-PikD and AVR-PikF) present in some isolates of the rice blast fungus, *Magnaporthe oryzae* with only AVR-PikD being recognised by the Pik resistance protein (Longya et al., 2019). It could be advantageous for the pathogen to have multiple *NIP2* and *NLP* genes with comparable functions expressed simultaneously during *in planta* growth. If one of these paralogs was recognized by a barley R protein and the pathogen evolved to mutate or eliminate that specific *NIP2* gene to escape recognition, then other paralogs could still function to promote pathogen virulence without triggering plant defence responses.

However, despite the clear duplication of members within the NIP2 family, the actual role they play in facilitating disease is unclear. Wevelsiep et al. (1991) characterised a protein they identified as NIP2 from *R. commune* and provided evidence that this protein induced necrosis in barley. However, our attempts to induce necrosis by infiltrating barley leaves with purified NIP2.1 was unsuccessful. We produced NIP2.1 in *E. coli* strain SHuffle® with CyDisCo which can be used to produce other active necrosis-inducing proteins from other pathogenic fungi. Such examples include the Tox proteins from *Parastagonospora nodorum* which were shown to be highly active post heterologous expression and able to induce cell death (Yu et al., 2022). Extensive biochemical tests confirmed the purity and the correct folding of the heterologously expressed NIP2.1, suggesting it was an active protein (Yu et al., 2022). In the original study of NIP2 by Wevelsiep et al. (1991), they showed that a protein identified as NIP2 induced necrosis on barley. However, the sequence of the infiltrated protein from this work has never been published, calling into question whether there is a mis-match between the activity observed in this assay and the identified protein. Additional research on *NIP2.1* was conducted by Kirsten et al. (2012) where they analysed the *NIP2.1* sequence of UK7 strain, which was one of the *R. commune* isolates tested by Wevelsiep et al. (Wevelsiep et al., 1991), UK7. However, in this study they did not investigate the ability of NIP2.1 of UK7 to cause cell death in barley leaves.

The inability of NIP2.1 to elicit plant cell death on barley leaves is supported by the experiment conducted by Zaffarano et al. (2008) who analysed the ability of different *Rhynchosporium* species to infect barley leaves. *R. commune* was the only species of *Rhynchosporium* that can infect and produce visible symptoms on barley leaves, while other *Rhynchosporium* species isolated from rye and triticale (*R. secalis*) and *Agropyron repens* (*R. agropyri*) were unable to cause visible symptoms on inoculated barley leaves (Zaffarano et al., 2008, 2011). However, *R*. *secalis* and *R*. *agropyri* also possessed the *NIP2.1* gene (Penselin et al., 2016) suggesting that *NIP2.1* was not specific to *R. commune* for inducing necrosis on barley leaves. Collectively, and combined with our data, these studies cast doubt over the role of NIP2.1 (and potentially other NIP2 proteins) to induce cell death in barley and to now consider other potential functions of this highly expressed effector during disease.

In conclusion, our study revealed the diversity of *NIP2* and *NLP* genes in the *R. commune* and its sister species, which likely originated from ancient duplications. RNA-sequencing and expression analyses consistently showed up-regulation of these genes during the early stages of infection suggesting their role in promoting infection whilst functional studies demonstrated that NIP2.1 does not induce necrosis as previously reported. These findings enhance our understanding of paralogous effectors in *R. commune*.

## Acknowledgements

We thank Haochen Wei for her technical assistance with the RNA-seq analysis and Arild Ranlym Arifin for his technical assistance with phylogenetic tree analysis. We acknowledge RSB plant services and NCRIS facility team for supporting our research.

## Data availability

Raw genome sequencing, raw RNA-Seq, genome assembly of WAI453, and the genome annotation of WAI453 are deposited under NCBI project PRJNA910335. Genome assembly and genome annotation of WAI453 are also available in Zenodo (DOI:10.5281/zenodo.13651118). All raw genome sequencing of other Australian_New isolates are deposited under NCBI project PRJNA914478.

Reviewer Link to PRJNA910335: https://dataview.ncbi.nlm.nih.gov/object/PRJNA910335?reviewer=2eiblqevmfsqradlia 7bu0l433

Reviewer Link to PRJNA914478: https://dataview.ncbi.nlm.nih.gov/object/PRJNA914478?reviewer=dq5crvah2dupkitcc 5p8r2ma7i

## CRediT authorship contribution statement

Reynaldi Darma: Writing – original draft, conceptualization, methodology, software, validation, formal analysis, investigation, data curation, visualization, funding acquisition

Daniel S. Yu: Writing- review & editing, conceptualization, methodology, formal analysis, investigation

Megan A. Outram: Writing- review & editing, conceptualization, methodology, formal analysis, investigation, visualization

Yi-Chang Sung: Writing- review & editing, methodology, investigation

Erin H. Hill: Writing- review & editing, methodology, software, data curation

Daniel Croll: Writing- review & editing, investigation

Simon J. Williams: Writing- review & editing, conceptualization, formal analysis, visualization

Ben Ovenden: Writing- review & editing, investigation

Andrew Milgate: Writing- review & editing, resources,

Peter S. Solomon: Writing- review & editing, conceptualization, methodology, supervision, project administration, funding acquisition

Megan C. McDonald: Writing- review & editing, conceptualization, methodology, supervision, project administration, funding acquisition

---

**Table S1.** PacBio sequencing result summary of *R. commune* WAI453 genome.

| Parameter | Value |
| --- | --- |
| Yield (bases) | 13,471,700,457 |
| Average read length (bases) | 11,390 |
| Longest read length (bases) | 90,133 |
| Reads used for genome assembly (>11kb)<br>(bases) | 10,614,786,115 |

**Table S2.** List of *Rhynchosporium* isolates used in this study.

| Species | Number of isolates |
| --- | --- |
| *R. agropyri* | 8 |
| *R. orthosporum* | 4 |
| *R. secalis* | 9 |
| *R. commune* | 190 |

**Table S3.**
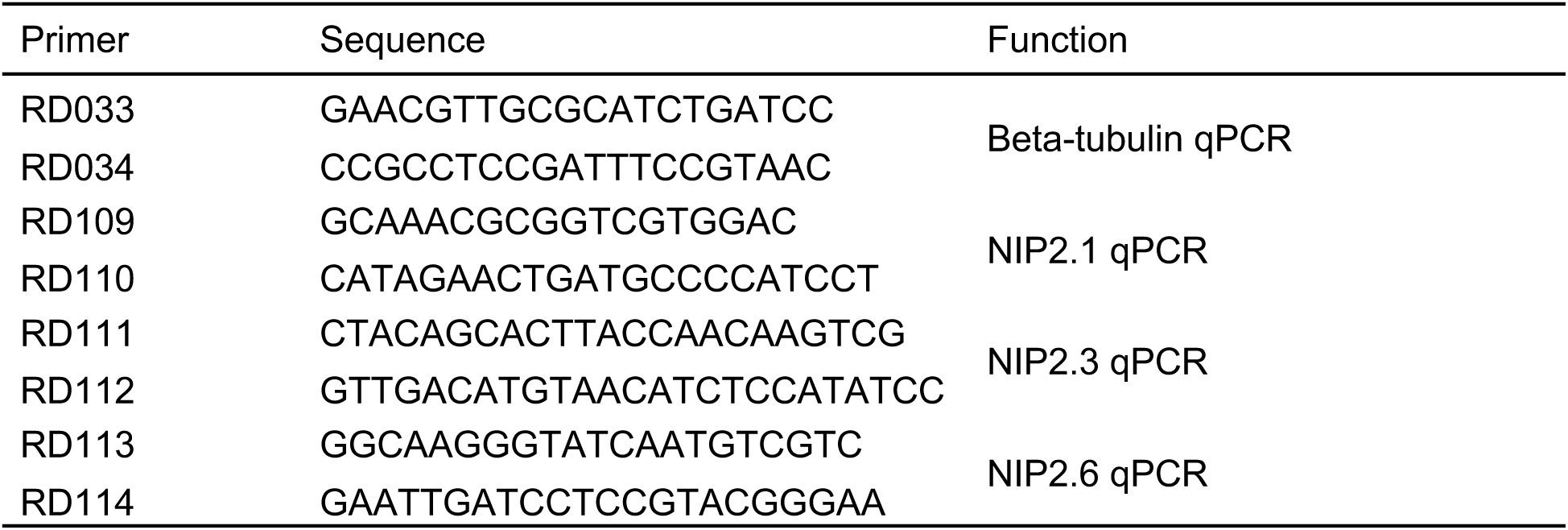
List of primer used in this study.

**Figure S1.**
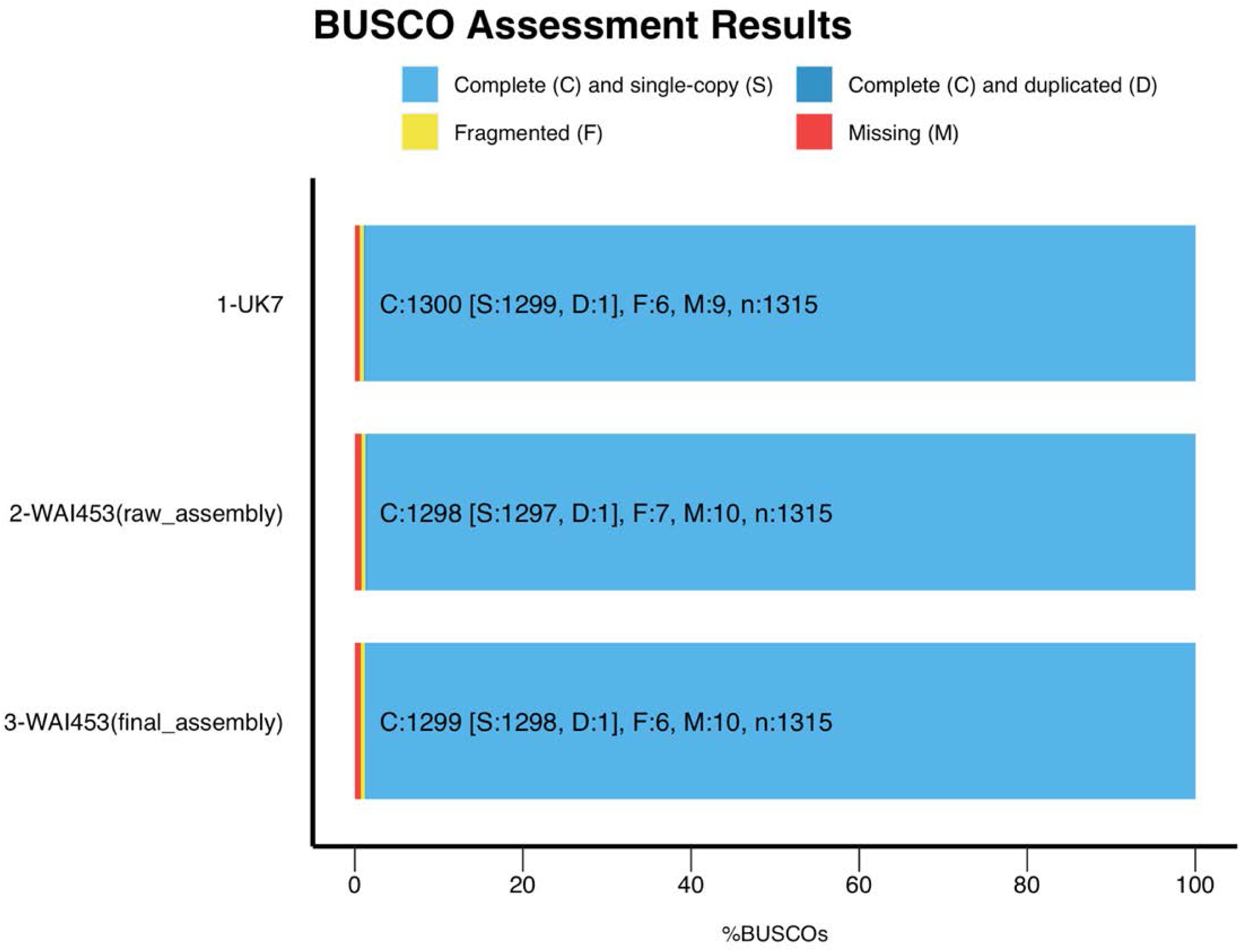
The completeness of the Ascomycota single copy orthologs genes (1,315 orthologs) from raw assembly of *R. commune* WAI453 and final assembly of *R. commune* WAI453 were assessed with the Benchmarking Universal Single-Copy Orthologs (BUSCO) tool. The *R. commune* UK7 genome was used as a control.

**Figure S2.**
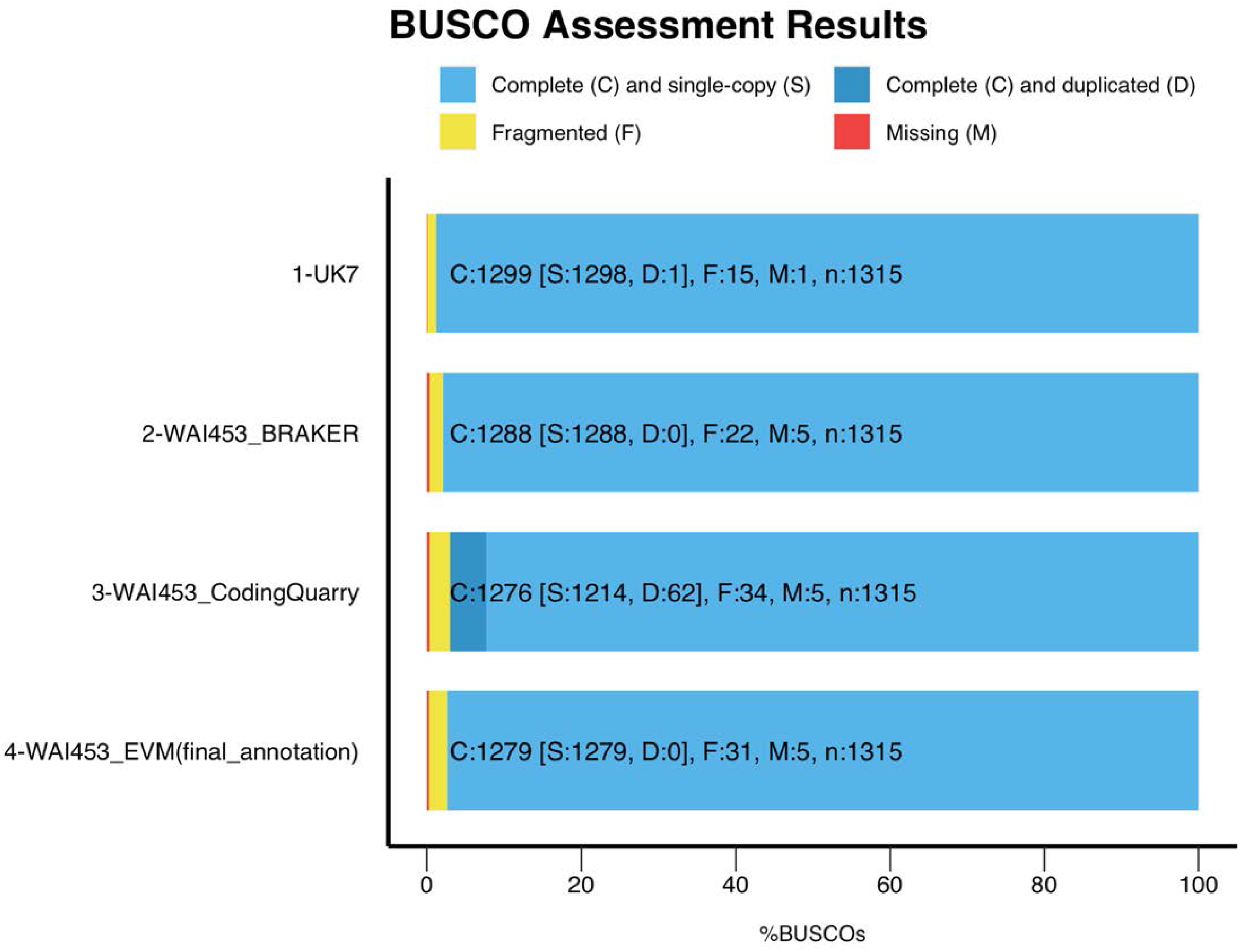
The completeness of the set of orthologs of the Ascomycota single copy genes (1,315 orthologs) from the raw genome annotation and the final genome annotation were assessed in BUSCO using the annotated amino acid sequences. The *R. commune* UK7 genome annotation was used as a control.

**Figure S3.**
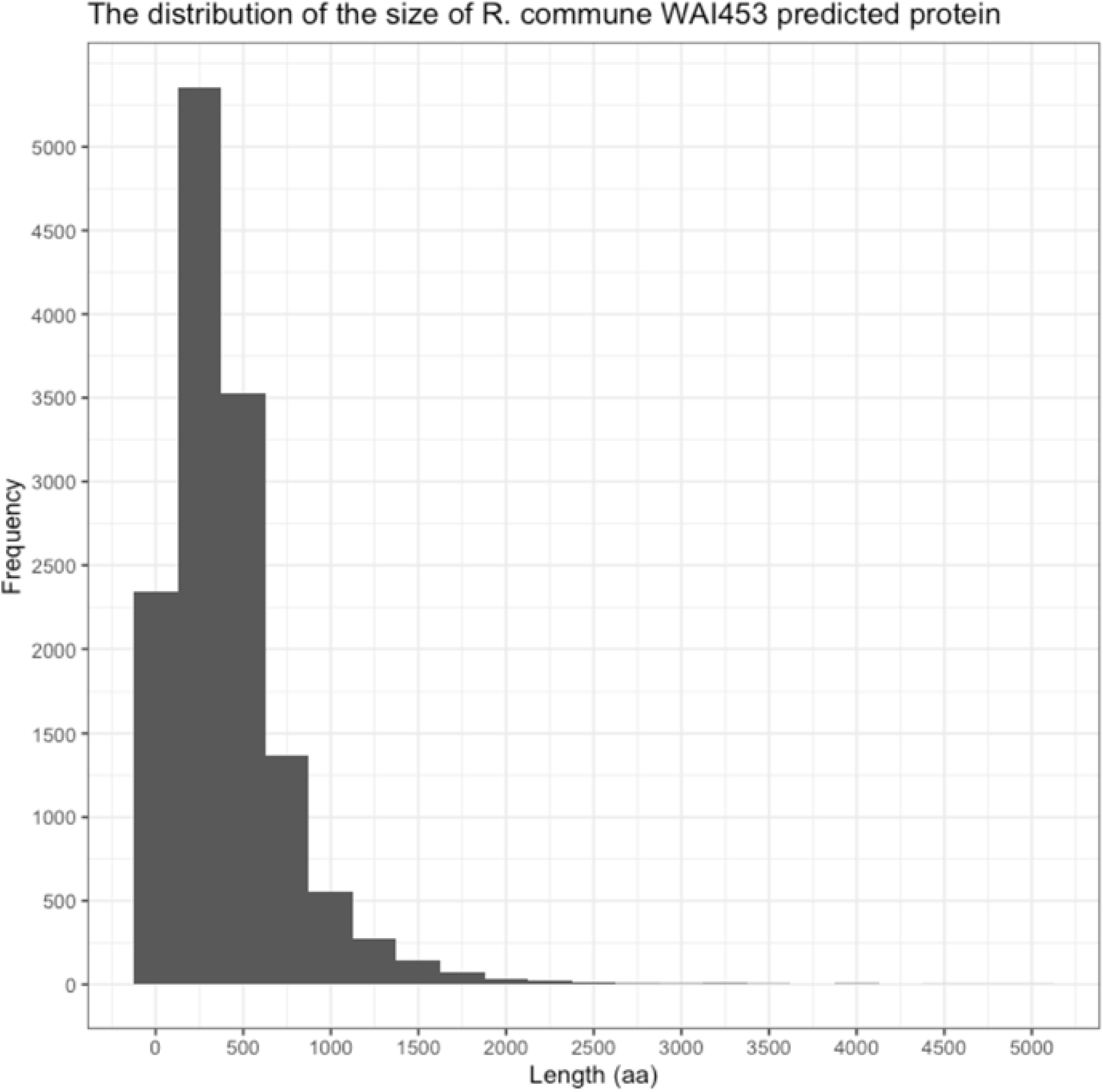
The size distribution of the *R. commune* WAI453 predicted proteins from the final annotation.

**Figure S4.**
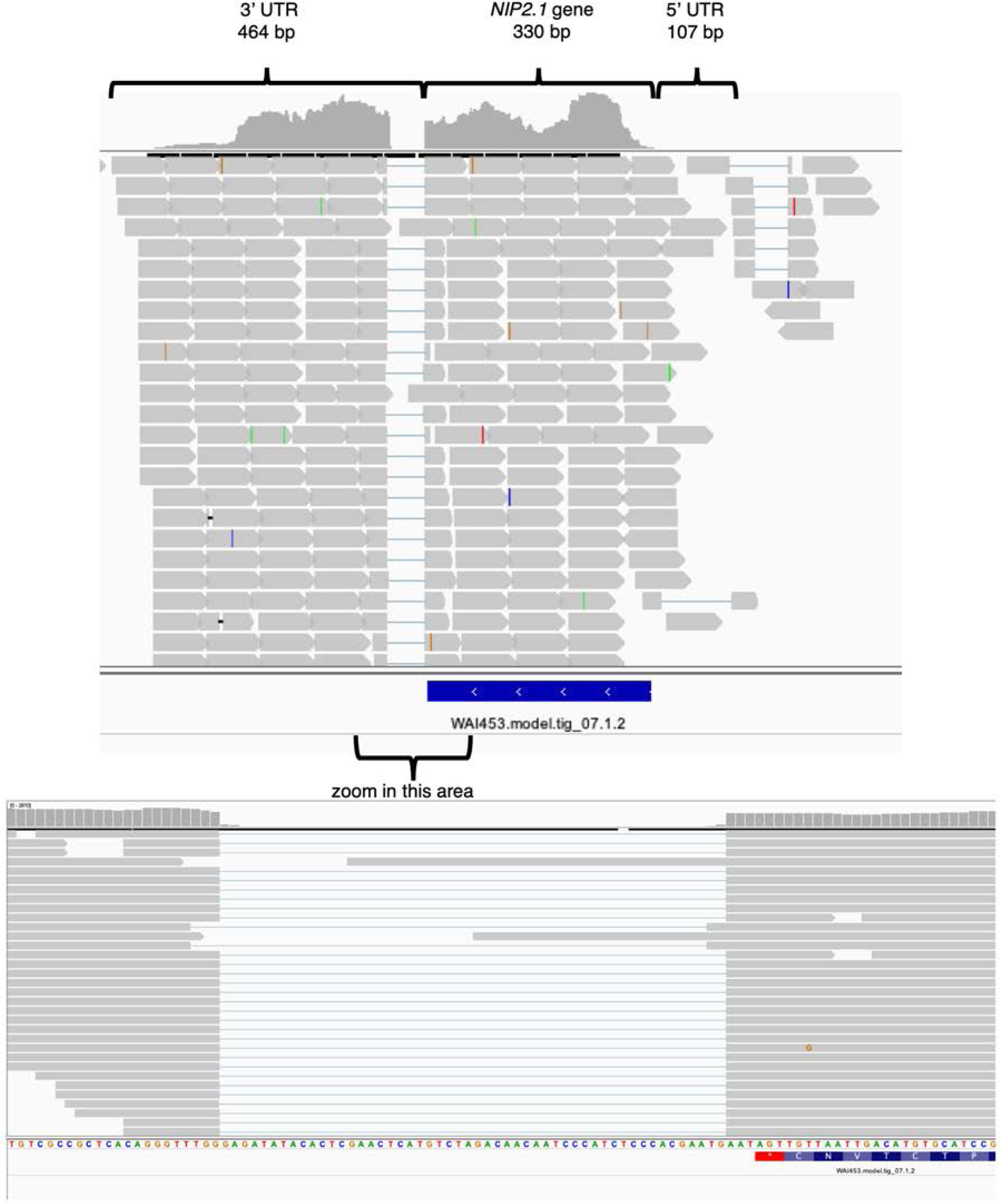
Visualisation of the mRNA reads mapped to *NIP2.1* region in the *R. commune* WAI453 genome in IGV viewer showed an additional intron after the stop codon in the *NIP2.1* transcript.

**Figure S5.**
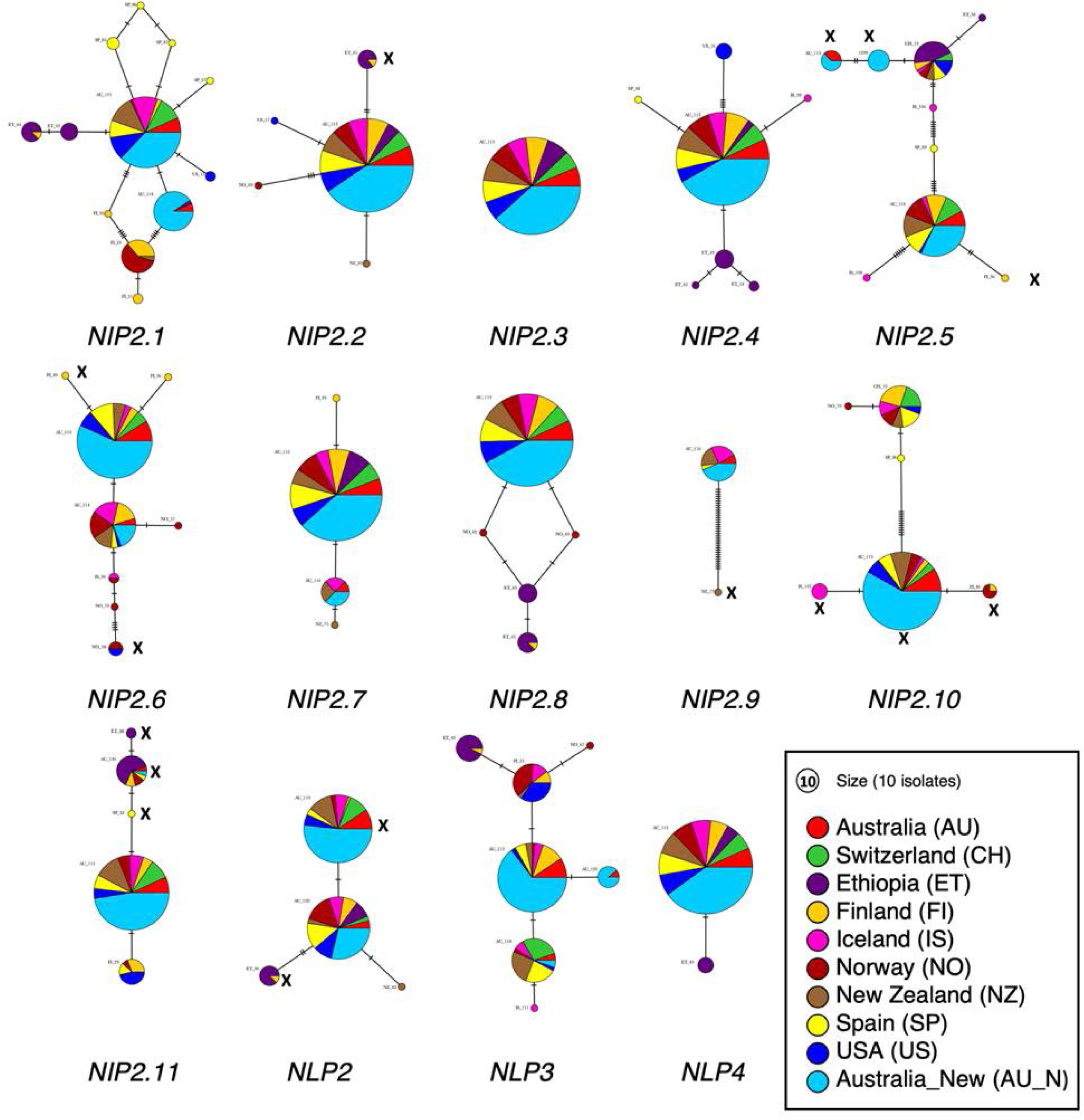
Haplotype networks of *NIP2* and *NLP* genes of *R. commune*. The haplotype networks were generated in the PopART with using minimum spanning network. *NIP2* genes with any mutation or any deletion and insertion are included in the analysis but any truncated genes are excluded in the analysis.

**Figure S6.**
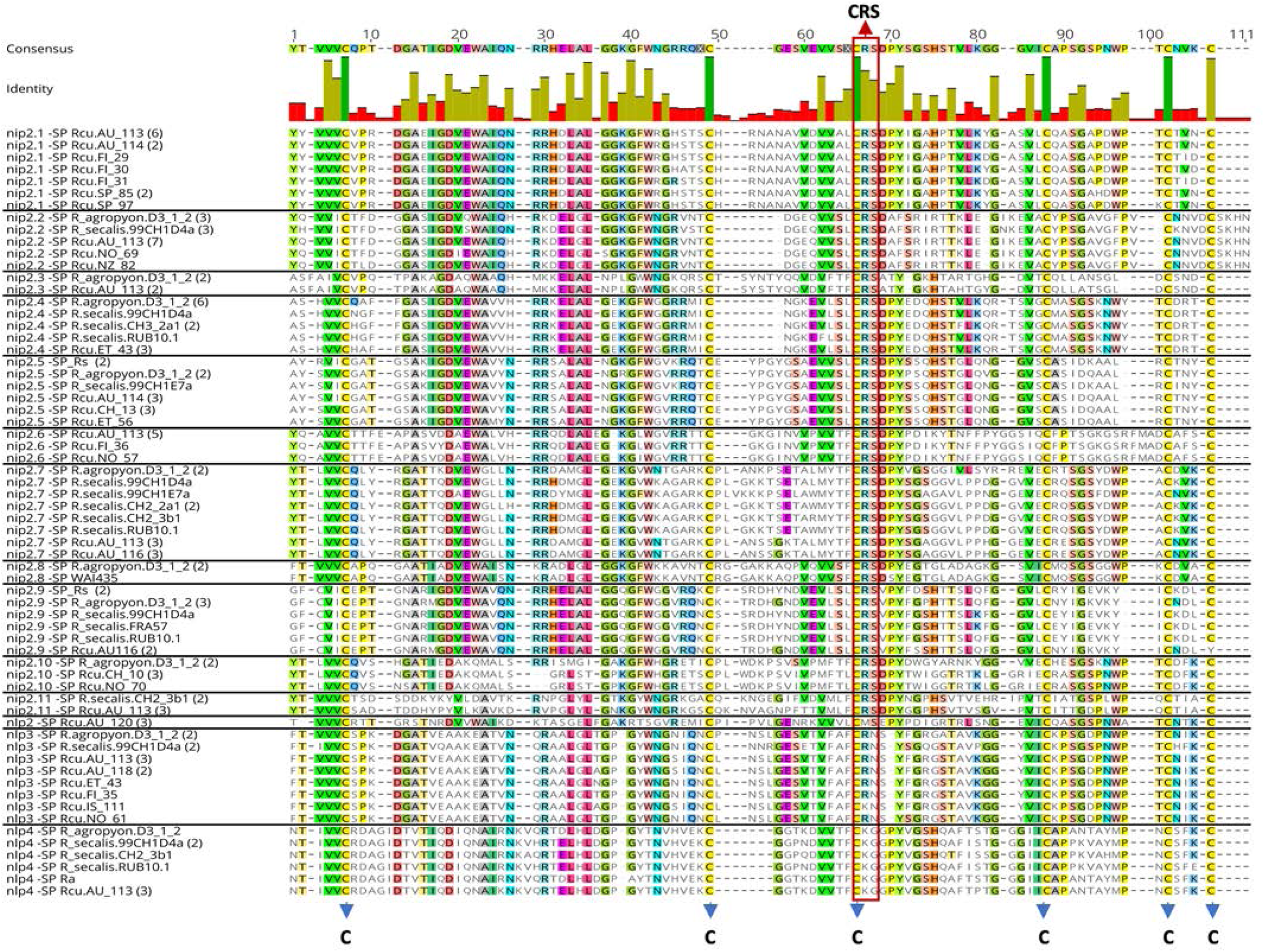
Alignment of 64 mature (without signal peptide) and complete (full length/without premature stop codon) NIP2 and NLP protein sequences from *R. commune* and the sister species. The conserved cysteine residues and the CRS domain are labelled with blue arrow and red arrow, respectively.

**Figure S7.**
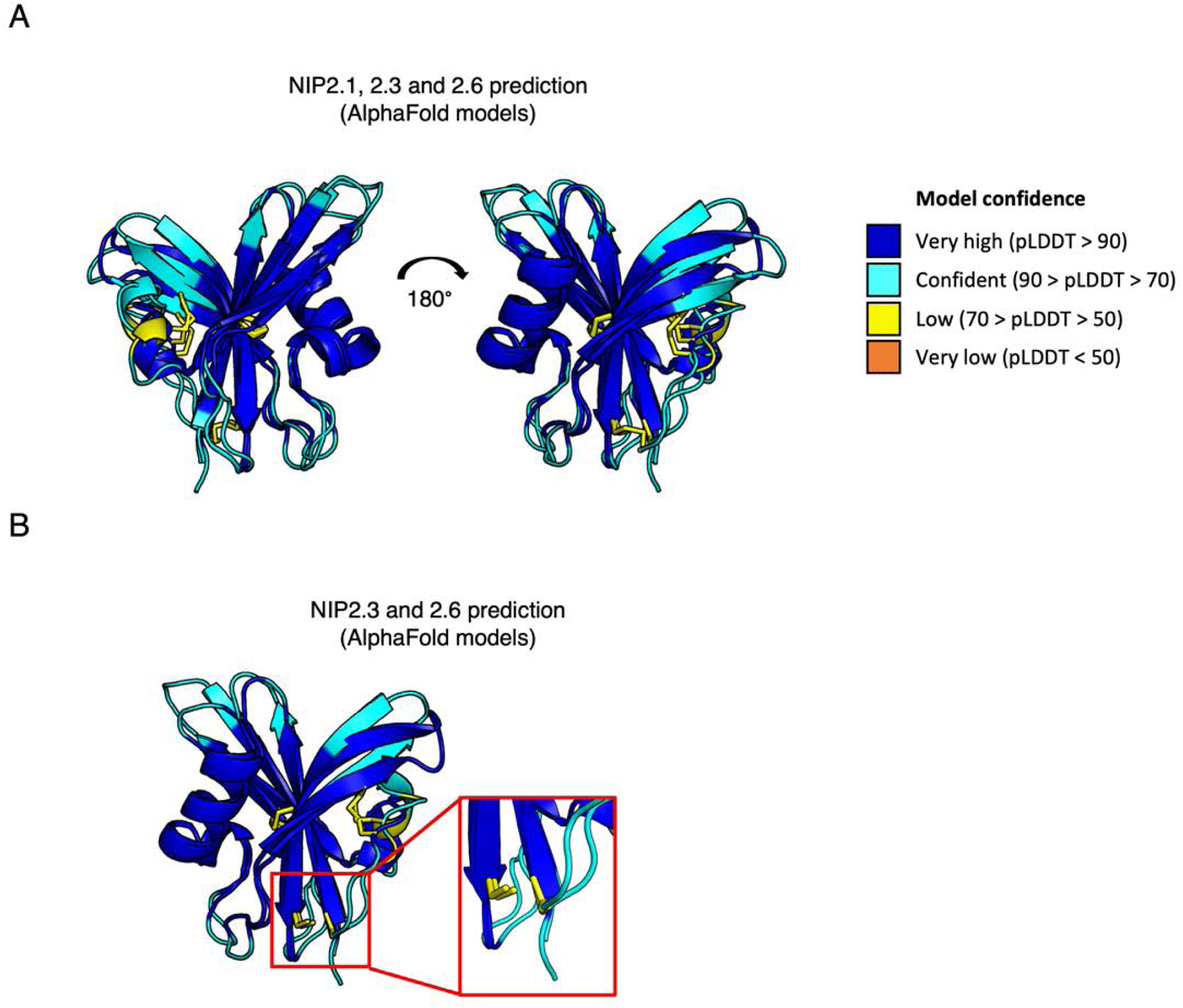
An AlphaFold 2 predictions of NIP2 proteins aligned (A) NIP2.1, NIP2.3, and NIP2.6 (B) NIP2.3 and NIP2.6, coloured according to pLDDT score as shown in the legend. NIP2.3 or NIP2.6 contains only two predicted disulphide bonds. The 3^rd^ site between residues 66 and 107 (Figure S6) were not predicted to form a disulfide based on the AlphaFold 2 prediction but those cysteines are conserved and localise to the same area with the 3^rd^ disulphide of NIP2.1. Cysteine residues or disulphide bonds shown as sticks and in yellow.

## References

Andrews, S., 2010.FastQC: a quality control tool for high throughput sequence data.

Avrova, A., Knogge, W., 2012. *Rhynchosporium commune*: a persistent threat to barley cultivation. Mol. Plant Pathol. 13, 986–997.

Bankevich, A., Nurk, S., Antipov, D., Gurevich, A.A., Dvorkin, M., Kulikov, A.S., Lesin, V.M., Nikolenko, S.I., Pham, S., Prjibelski, A.D., Pyshkin, A.V., Sirotkin, A.V., Vyahhi, N., Tesler, G., Alekseyev, M.A., Pevzner, P.A., 2012. SPAdes: a new genome assembly algorithm and its applications to single-cell sequencing. J. Comput. Biol. 19, 455–477.

Benjamini, Y., Hochberg, Y., 1995. Controlling the false discovery rate: A practical and powerful approach to multiple testing. J. R. Stat. Soc. Ser. B Stat. Methodol. 57, 289–300.

Bolger, A.M., Lohse, M., Usadel, B., 2014. Trimmomatic: a flexible trimmer for Illumina sequence data. Bioinformatics 30, 2114–2120.

Brown, J.S., 1985. Pathogenic variation among isolates of *Rhynchosporium secalis* from cultivated barley growing in Victoria, Australia. Euphytica 34, 129–133.

Brůna, T., Hoff, K.J., Lomsadze, A., Stanke, M., Borodovsky, M., 2021.BRAKER2: automatic eukaryotic genome annotation with GeneMark-EP+ and AUGUSTUS supported by a protein database. NAR Genom. Bioinform. 3, lqaa108.

Camacho, C., Coulouris, G., Avagyan, V., Ma, N., Papadopoulos, J., Bealer, K., Madden, T.L., 2009. BLAST+: architecture and applications. BMC Bioinformatics 10, 421.

Chen, L.H., Kracun, S.K., Nissen, K.S., Mravec, J., Jorgensen, B., Labavitch, J., Stergiopoulos, I., 2021. A diverse member of the fungal Avr4 effector family interacts with de-esterified pectin in plant cell walls to disrupt their integrity. Sci. Adv. 7, eabe0809.

Dobin, A., Davis, C.A., Schlesinger, F., Drenkow, J., Zaleski, C., Jha, S., Batut, P., Chaisson, M., Gingeras, T.R., 2013. STAR: ultrafast universal RNA-seq aligner. Bioinformatics 29, 15–21.

Fiegen, M., Knogge, W., 2002. Amino acid alterations in isoforms of the effector protein NIP1 from *Rhynchosporium secalis* have similar effects on its avirulence- and virulence-associated activities on barley. Physiol. Mol. Plant Pathol. 61, 299–302.

Gardiner, D.M., Cozijnsen, A.J., Wilson, L.M., Pedras, M.S., Howlett, B.J., 2004. The sirodesmin biosynthetic gene cluster of the plant pathogenic fungus *Leptosphaeria maculans*. Mol. Microbiol. 53, 1307–1318.

Goodwin, S.B., 2002. The barley scald pathogen *Rhynchosporium secalis* is closely related to the discomycetes *Tapesia* and *Pyrenopeziza*. Mycol. Res. 106, 645–654.

Goodwin, S.B., Allard, R.W., Webster, R.K., 1990. A nomenclature for *Rhynchosporium secalis* pathotypes. Phytopathology 80, 1330–1336.

Haas, B.J., Salzberg, S.L., Zhu, W., Pertea, M., Allen, J.E., Orvis, J., White, O., Buell, C.R., Wortman, J.R., 2008. Automated eukaryotic gene structure annotation using EVidenceModeler and the program to assemble spliced alignments. Genome Biol. 9, R7.

Hatahet, F., Nguyen, V.D., Salo, K.E.H., Ruddock, L.W., 2010. Disruption of reducing pathways is not essential for efficient disulfide bond formation in the cytoplasm of *E. coli*. Microb. Cell Fact. 9,67.

Holm, L., Rosenstrom, P., 2010. Dali server: conservation mapping in 3D. Nucleic Acids Res. 38, W545–W549.

Huelsenbeck, J.P., Ronquist, F., 2001. MRBAYES: Bayesian inference of phylogenetic trees. Bioinformatics 17, 754–755.

Jumper, J., Evans, R., Pritzel, A., Green, T., Figurnov, M., Ronneberger, O., Tunyasuvunakool, K., Bates, R., Zidek, A., Potapenko, A., Bridgland, A., Meyer, C., Kohl, S.A.A., Ballard, A.J., Cowie, A., Romera-Paredes, B., Nikolov, S., Jain, R., Adler, J., Back, T., Petersen, S., Reiman, D., Clancy, E., Zielinski, M., Steinegger, M., Pacholska, M., Berghammer, T., Bodenstein, S., Silver, D., Vinyals, O., Senior, A.W., Kavukcuoglu, K., Kohli, P., Hassabis, D., 2021. Highly accurate protein structure prediction with AlphaFold. Nature 596, 583–589.

Kalyaanamoorthy, S., Minh, B.Q., Wong, T.K.F., von Haeseler, A., Jermiin, L.S., 2017. ModelFinder: fast model selection for accurate phylogenetic estimates. Nat. Methods 14, 587–589.

Kearse, M., Moir, R., Wilson, A., Stones-Havas, S., Cheung, M., Sturrock, S., Buxton, S., Cooper, A., Markowitz, S., Duran, C., Thierer, T., Ashton, B., Meintjes, P., Drummond, A., 2012. Geneious Basic: an integrated and extendable desktop software platform for the organization and analysis of sequence data. Bioinformatics 28, 1647–1649.

King, K.M., West, J.S., Brunner, P.C., Dyer, P.S., Fitt, B.D.L., 2013. Evolutionary relationships between *Rhynchosporium lolii* sp nov and other *Rhynchosporium* species on grasses. PLoS One 8, e72536.

Kirsten, S., Navarro-Quezada, A., Penselin, D., Wenzel, C., Matern, A., Leitner, A., Baum, T., Seiffert, U., Knogge, W., 2012. Necrosis-inducing proteins of *Rhynchosporium commune*, effectors in quantitative disease resistance. Mol. Plant Microbe Interact. 25, 1314–1325.

Koren, S., Walenz, B.P., Berlin, K., Miller, J.R., Bergman, N.H., Phillippy, A.M., 2017. Canu: scalable and accurate long-read assembly via adaptive k-mer weighting and repeat separation. Genome Res. 27, 722–736.

Leigh, J.W., Bryant, D., 2015. PopART: Full-feature software for haplotype network construction. Methods Ecol. Evol. 6, 1110–1116.

Linde, C.C., Zala, M., Ceccarelli, S., McDonald, B.A., 2003. Further evidence for sexual reproduction in *Rhynchosporium secalis* based on distribution and frequency of mating-type alleles. Fungal Genet. Biol. 40, 115–125.

Linde, C.C., Zala, M., McDonald, B.A., 2009. Molecular evidence for recent founder populations and human-mediated migration in the barley scald pathogen *Rhynchosporium secalis*. Mol. Phylogenet. Evol. 51, 454–464.

Lobstein, J., Emrich, C.A., Jeans, C., Faulkner, M., Riggs, P., Berkmen, M., 2016. Erratum to: SHuffle, a novel *Escherichia coli* protein expression strain capable of correctly folding disulfide bonded proteins in its cytoplasm. Microb. Cell Fact. 15, 124.

Longya, A., Chaipanya, C., Franceschetti, M., Maidment, J.H.R., Banfield, M.J., Jantasuriyarat, C., 2019. Gene duplication and mutation in the emergence of a novel aggressive allele of the *AVR-Pik* effector in the rice blast fungus. Mol. Plant Microbe Interact. 32, 740–749.

Lu, X., Miao, J., Shen, D., Dou, D., 2022. Proteinaceous effector discovery and characterization in plant pathogenic *Colletotrichum* fungi. Front. Microbiol. 13, 914035.

Lynch, M., Conery, J.S., 2000. The evolutionary fate and consequences of duplicate genes. Science 290, 1151–1155.

Matos, C.F.R.O., Robinson, C., Alanen, H.I., Prus, P., Uchida, Y., Ruddock, L.W., Freedman, R.B., Keshavarz-Moore, E., 2014. Efficient export of prefolded, disulfide-bonded recombinant proteins to the periplasm by the Tat pathway in *Escherichia coli* CyDisCo strains. Biotechnol. Prog. 30, 281–290.

McDermott, J.M., McDonald, B.A., Allard, R.W., Webster, R.K., 1989. Genetic variability for pathogenicity, isozyme, ribosomal DNA and colony color variants in populations of *Rhynchosporium secalis*. Genetics 122, 561–565.

McDonald, B.A., Zhan, J., Burdon, J.J., 1999. Genetic structure of *Rhynchosporium secalis* in Australia. Phytopathology 89, 639–645.

McDonald, M.C., Ahren, D., Simpfendorfer, S., Milgate, A., Solomon, P.S., 2018. The discovery of the virulence gene *ToxA* in the wheat and barley pathogen *Bipolaris sorokiniana*. Mol. Plant Pathol. 19, 432–439.

Mirdita, M., Schütze, K., Moriwaki, Y., Heo, L., Ovchinnikov, S., Steinegger, M., 2022. ColabFold: making protein folding accessible to all. Nat. Methods 19, 679–682.

Mohd-Assaad, N., McDonald, B.A., Croll, D., 2019. The emergence of the multi-species NIP1 effector in *Rhynchosporium* was accompanied by high rates of gene duplications and losses. Environ. Microbiol 21, 2677–2695.

Muller, P.Y., Janovjak, H., Miserez, A.R., Dobbie, Z., 2002. Processing of gene expression data generated by quantitative real-time RT-PCR. Biotechniques 32, 1372–1374, 1376, 1378-1379.

Navarrete, F., Grujic, N., Stirnberg, A., Saado, I., Aleksza, D., Gallei, M., Adi, H., Alcantara, A., Khan, M., Bindics, J., Trujillo, M., Djamei, A., 2021. The Pleiades are a cluster of fungal effectors that inhibit host defenses. PLoS Pathog. 17, e1009641.

Nguyen, L.T., Schmidt, H.A., von Haeseler, A., Minh, B.Q., 2015. IQ-TREE: a fast and effective stochastic algorithm for estimating maximum-likelihood phylogenies. Mol. Biol. Evol. 32, 268–274.

Outram, M.A., Sung, Y.C., Yu, D., Dagvadorj, B., Rima, S.A., Jones, D.A., Ericsson, D.J., Sperschneider, J., Solomon, P.S., Kobe, B., Williams, S.J., 2021. The crystal structure of SnTox3 from the necrotrophic fungus *Parastagonospora nodorum* reveals a unique effector fold and provides insight into Snn3 recognition and pro-domain protease processing of fungal effectors. New Phytol. 231, 2282–2296.

Paveley, N., Fitt, B., Oxley, S.J.P., Bingham, I.J., Cockerell, V., Edwars, C., Dodgson, G., Gosling, P., Nicholls, C., Watts, J., Boys, E., Geary, F., 2016. Barley disease management guide, AHDB (ed) Agriculture and Horticulture Development Board Cereals & Oilseeds, Warwickshire.

Penselin, D., Munsterkotter, M., Kirsten, S., Felder, M., Taudien, S., Platzer, M., Ashelford, K., Paskiewicz, K.H., Harrison, R.J., Hughes, D.J., Wolf, T., Shelest, E., Graap, J., Hoffmann, J., Wenzel, C., Woltje, N., King, K.M., Fitt, B.D., Guldener, U., Avrova, A., Knogge, W., 2016. Comparative genomics to explore phylogenetic relationship, cryptic sexual potential and host specificity of *Rhynchosporium* species on grasses. BMC Genomics 17, 953.

Pertea, M., Pertea, G.M., Antonescu, C.M., Chang, T.C., Mendell, J.T., Salzberg, S.L., 2015. StringTie enables improved reconstruction of a transcriptome from RNA-seq reads. Nat. Biotechnol 33, 290–295.

Rambaut, A., 2018. FigTree v1.4.4: Molecular Evolution, Phylogenetics and Epidemiology. University of Edinburgh, Edinburgh.

Ridout, C.J., Skamnioti, P., Porritt, O., Sacristan, S., Jones, J.D., Brown, J.K., 2006. Multiple avirulence paralogues in cereal powdery mildew fungi may contribute to parasite fitness and defeat of plant resistance. Plant Cell 18, 2402–2414.

Robinson, M.D., McCarthy, D.J., Smyth, G.K., 2010. edgeR: a Bioconductor package for differential expression analysis of digital gene expression data. Bioinformatics 26, 139–140.

Salamati, S., Zhan, J., Burdon, J.J., McDonald, B.A., 2000. The genetic structure of field populations of *Rhynchosporium secalis* from three continents suggests moderate gene flow and regular recombination. Phytopathology 90, 901–908.

Saur, I.M., Bauer, S., Kracher, B., Lu, X., Franzeskakis, L., Muller, M.C., Sabelleck, B., Kummel, F., Panstruga, R., Maekawa, T., Schulze-Lefert, P., 2019. Multiple pairs of allelic MLA immune receptor-powdery mildew AVR_A_ effectors argue for a direct recognition mechanism. Elife 8, e44471.

Seong, K., Krasileva, K.V., 2023. Prediction of effector protein structures from fungal phytopathogens enables evolutionary analyses. Nat. Microbiol 8, 174–187.

Shao, D., Smith, D.L., Kabbage, M., Roth, M.G., 2021. Effectors of plant necrotrophic fungi. Front. Plant Sci. 12, 687713.

Shen, D., Liu, T., Ye, W., Liu, L., Liu, P., Wu, Y., Wang, Y., Dou, D., 2013. Gene duplication and fragment recombination drive functional diversification of a superfamily of cytoplasmic effectors in *Phytophthora sojae*. PLoS One 8, e70036.

Simao, F.A., Waterhouse, R.M., Ioannidis, P., Kriventseva, E.V., Zdobnov, E.M., 2015. BUSCO: assessing genome assembly and annotation completeness with single-copy orthologs. Bioinformatics 31, 3210–3212.

Skoropad, W.P., 1959. Seed and seedling infection of barley by *Rhynchosporium secalis*. Phytopathology 49, 623–626.

Stamatakis, A., 2014. RAxML version 8: a tool for phylogenetic analysis and post-analysis of large phylogenies. Bioinformatics 30, 1312–1313.

Stefansson, T.S., McDonald, B.A., Willi, Y., 2013. Local adaptation and evolutionary potential along a temperature gradient in the fungal pathogen *Rhynchosporium commune*. Evol. Appl. 6, 524–534.

Testa, A.C., Hane, J.K., Ellwood, S.R., Oliver, R.P., 2015. CodingQuarry: highly accurate hidden Markov model gene prediction in fungal genomes using RNA-seq transcripts. BMC Genomics 16, 170.

van Kempen, M., Kim, S.S., Tumescheit, C., Mirdita, M., Lee, J., Gilchrist, C.L.M., Soding, J., Steinegger, M., 2023. Fast and accurate protein structure search with Foldseek. Nat. Biotechnol. 42, 243–246.

Wallwork, H., Grcic, M., 2011. The use of differential isolates of *Rhynchosporium secalis* to identify resistance to leaf scald in barley. Austral. Plant Pathol. 40, 490–496.

Wevelsiep, L., Kogel, K.-H., Knogge, W., 1991. Purification and characterization of peptides from *Rhynchosporium secalis* inducing necrosis in barley. Physiol. Mol. Plant Pathol. 39, 471–482.

Wevelsiep, L., Rupping, E., Knogge, W., 1993. Stimulation of barley plasmalemma H+-ATPase by phytotoxic peptides from the fungal pathogen *Rhynchosporium secalis*. Plant Physiol. 101, 297–301.

Yu, D.S., Outram, M.A., Crean, E., Smith, A., Sung, Y.C., Darma, R., Sun, X., Ma, L., Jones, D.A., Solomon, P.S., Williams, S.J., 2022. Optimised production of disulfide-bonded fungal effectors in *E. coli* using CyDisCo and FunCyDisCo co-expression approaches. Mol. Plant Microbe Interact. 35, 109–118.

Yu, D.S., Outram, M.A., Smith, A., McCombe, C.L., Khambalkar, P.B., Rima, S.A., Sun, X., Ma, L., Ericsson, D.J., Jones, D.A., Williams, S.J., 2024. The structural repertoire of *Fusarium oxysporum* f. sp. *lycopersici* effectors revealed by experimental and computational studies. Elife 12, RP89280.

Zaffarano, P.L., McDonald, B.A., Linde, C.C., 2008. Rapid speciation following recent host shifts in the plant pathogenic fungus *Rhynchosporium*. Evolution 62, 1418–1436.

Zaffarano, P.L., McDonald, B.A., Linde, C.C., 2011. Two new species of *Rhynchosporium*. Mycologia 103, 195–202.

Zhan, J., Fitt, B.D.L., Pinnschmidt, H.O., Oxley, S.J.P., Newton, A.C., 2008. Resistance, epidemiology and sustainable management of *Rhynchosporium secalis* populations on barley. Plant Pathol. 57, 1–14.

Zhang, X., Nguyen, N., Breen, S., Outram, M.A., Dodds, P.N., Kobe, B., Solomon, P.S., Williams, S.J., 2016. Production of small cysteine-rich effector proteins in *Escherichia coli* for structural and functional studies. Mol. Plant Pathol. 18, 141–151.

Zhang, X., Ovenden, B., Milgate, A., 2020. Recent insights into barley and *Rhynchosporium commune* interactions. Mol. Plant Pathol. 21, 1111–1128.

